# Neuron-specific increase in lamin B1 disrupts nuclear function in Huntington’s disease

**DOI:** 10.1101/2020.03.06.979674

**Authors:** Rafael Alcalá-Vida, Marta Garcia-Forn, Jordi Creus-Muncunill, Yoko Ito, Enrique Blanco, Carla Castany-Pladevall, Killian Crespí-Vázquez, Aled Parry, Guy Slater, Shamith Samarajiwa, Sandra Peiró, Luciano Di Croce, Masashi Narita, Esther Pérez-Navarro

**Affiliations:** Departament de Biomedicina, Facultat de Medicina i Ciències de la Salut, Institut de Neurociències, Universitat de Barcelona, 08036 Barcelona, Catalonia; Institut d’Investigacions Biomèdiques August Pi i Sunyer (IDIBAPS), Barcelona, Catalonia; Centro de Investigación Biomédica en Red sobre Enfermedades Neurodegenerativas (CIBERNED), Spain; Cancer Research UK Cambridge Institute, University of Cambridge, Cambridge, UK; Centre for Genomic Regulation (CRG), The Barcelona Institute of Science and Technology, Dr. Aiguader 88, Barcelona, Spain; Epigenetics Programme, The Babraham Institute, Cambridge, UK; MRC Cancer Unit, Hutchison/MRC Research Centre, University of Cambridge, Cambridge Biomedical Campus, Cambridge, CB2 0XZ, UK; Vall d’Hebron Institute of Oncology, 08035, Barcelona, Spain; ICREA, Pg. Lluis Companys 23, Barcelona, Spain; Actual address: Laboratory of Cognitive and Adaptive Neuroscience, UMR 7364 (CNRS/Strasbourg University), Strasbourg, France

**Keywords:** chromatin accessibility, LAD, R6/1 mouse, nuclear morphology, nuclear permeability

## Abstract

Lamins are crucial proteins for nuclear functionality. Here, we provide new evidence showing an involvement of increased lamin B1 levels in the pathophysiology of Huntington’s disease (HD), a CAG repeat-associated neurodegenerative disorder. Through fluorescence-activated nuclear suspension imaging we demonstrate that nucleus from striatal medium-sized spiny and CA1 hippocampal neurons display increased lamin B1 levels, in correlation with altered nuclear morphology and nucleocytoplasmic transport disruption. Moreover, ChIP-sequencing analysis shows an alteration of lamin-associated chromatin domains in hippocampal nuclei, which could contribute to transcriptional alterations we determined by RNA sequencing. Supporting lamin B1 alterations as a causal role in mutant-huntingtin mediated neurodegeneration, pharmacological normalization of lamin B1 levels by betulinic acid administration in the R6/1 mouse model of HD restored nuclear homeostasis and prevented motor and cognitive dysfunction. Collectively, our work point out increased lamin B1 levels as a new pathogenic mechanism in HD and provides a novel target for its intervention.

## Introduction

Lamins are type V intermediate filaments that together with lamin-binding proteins are embedded into the inner nuclear membrane and constitute the nuclear lamina^1^. This family of proteins is classified in two subgroups: A-type lamins (lamins A and C), encoded by the *LMNA* gene, and B-type lamins (lamins B1 and B2), encoded by *LMNB1* and *LMNB2* genes, respectively^1^. It was long thought that their only function was to provide a structural support to the nuclear envelope membrane, but evidence indicates that they are involved in a wide variety of cell functions and processes, including DNA replication, transcription, chromatin organization and nucleus-cytoplasm interaction^2^. While lamins A and C are expressed exclusively in differentiated cells, lamin B is present in almost all cell types independently of their differentiation state^3^. This suggests that B-type lamins are essential for the survival of mammalian cells^4^.

Alterations in lamins content or structure leads to a particular type of nuclear envelopathies called laminopathies^5^. While many laminopathies are associated with mutations in *LMNA* gene^5^, only two have been associated with alterations in lamin B: the autosomal dominant leukodystrophy^6^ and the acquired partial lipodystrophy^7^ caused by *LMNB1* and *LMNB2* mutations, respectively. Interestingly, in the last few years, lamin B alterations have also been found in neurodegenerative disorders such as Parkinson’s (PD) and Alzheimer’s diseases^8,9^. Strikingly, two cellular functions in which lamin B plays a critical role, RNA nuclear exportation^10^ and nuclear pore complex organization^11^, are altered in Huntington’s disease (HD), an autosomal dominant neurodegenerative disorder caused by an inherited CAG repeat expansion in the exon-1 of the huntingtin (*htt*) gene^12^. This mutation results in the lengthening of the polyglutamine chain at the amino terminus of the huntingtin (Htt) protein inducing self-association and aggregation. Consequently, mutant Htt (mHtt) loses its biological functions and becomes toxic^13^. In HD, medium-sized spiny neurons (MSNs), the GABAergic output projection neurons that account for the vast majority (90-95%) of all striatal neurons, are mainly affected. Although motor symptoms are the most prominent, psychiatric alterations and cognitive decline appear first in HD patients, which become more evident as the disease progresses. Cognitive deficits are related to the dysfunction of the corticostriatal pathway and the hippocampus and, together with motor deficits, have been replicated in most HD mouse models^14^.

Molecular mechanisms leading to nuclear lamina alterations in neurons expressing mHtt remain to be elucidated. Previous results from our lab suggested that decreased levels of the pro-apoptotic kinase PKCδ would lead to an aberrant accumulation of lamin B^15^ which, in turn, could have a significant influence in the nuclear lamina structure^16,17^. Therefore, here we sought to deeply characterize the impact of lamin alterations in HD brain at physiological (studying nuclear lamina morphology and nucleo-cytoplasmic transport), transcriptomic (by generating RNA-sequencing (RNA-seq) data) and epigenetic (analyzing lamin chromatin binding and chromatin accessibility) levels by using the R6/1 transgenic mouse model of HD and human post-mortem brain samples.

## Results

### Lamin B levels are increased in a region-specific manner in HD brain

Lamin B1, B2, and A/C protein levels were analyzed in the striatum, cortex and hippocampus of wild-type and R6/1 mice, a transgenic mouse model of HD over-expressing the exon 1 of the human mutant huntingtin^18^, at different ages. Western blot analysis revealed an increase in lamin B1 and B2 levels in all three regions from early disease stages in R6/1 mice (Fig. 1a, b) whereas lamin A/C protein levelsremained unchanged until 30 weeks of age in the striatum and hippocampus (Supplementary Fig. 1). Since the most important alterations were found in lamin B isoforms, we investigated whether such an increase was reproducible in the brain of HD patients. Western blot analysis revealed that lamin B1 levels were significantly higher, in comparison to levels in non-affected individuals, in the putamen of HD patients at Vonsattel (VS) grade III-IV and in the frontal cortex of HD patients at grade I-II and III-IV (Fig. 1c). Unexpectedly, no significant changes were found within the hippocampus of HD patients at any disease stage. On the other hand, lamin B2 protein levels were only increased in the frontal cortex of HD patients (Fig. 1d). Consequently, only lamin B1 levels are consistently affected in the brain of R6/1 mice and HD patients.

**Figure 1.**
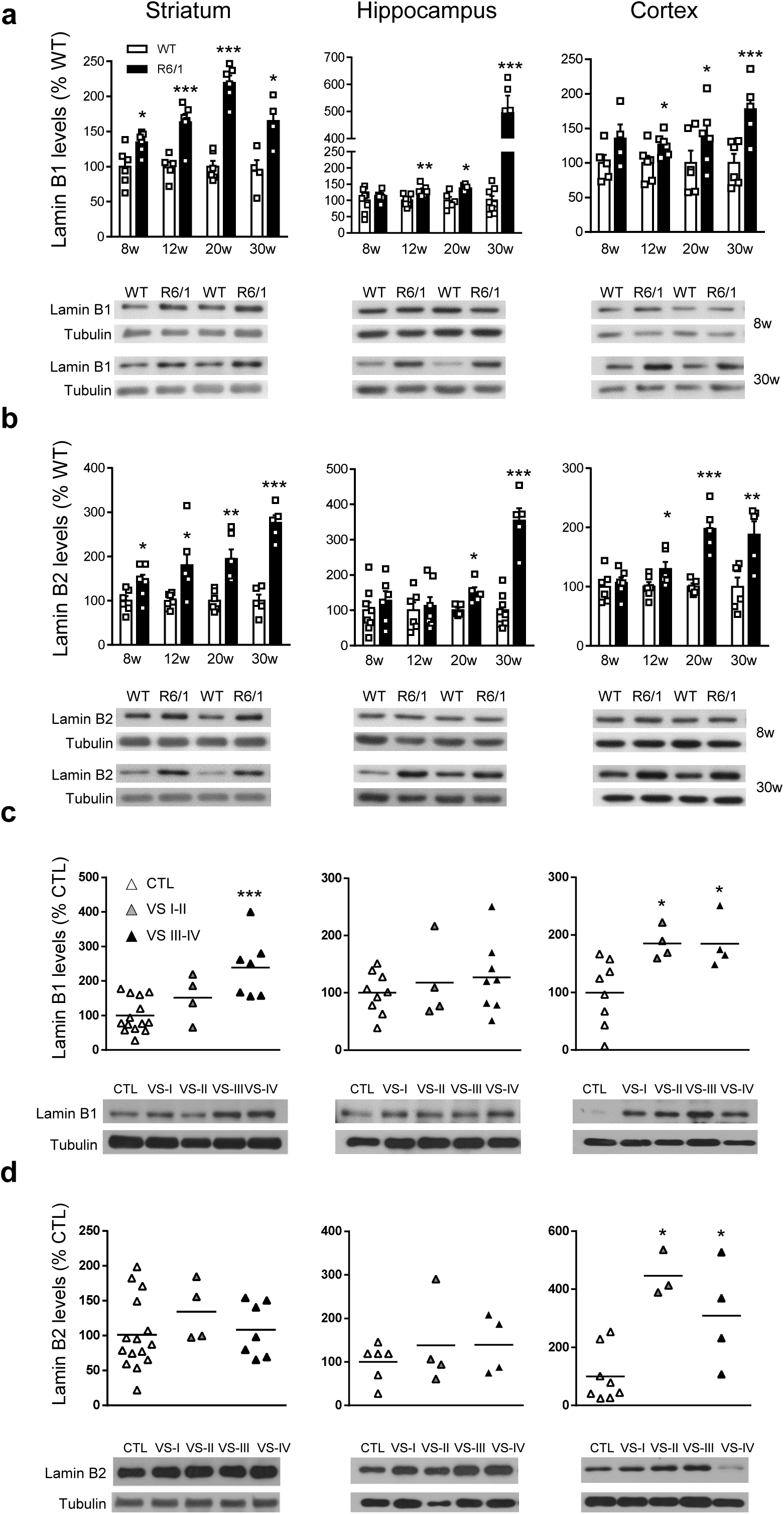
Lamin B1 and B2 are differentially increased in Huntington’s disease brain. (**a, c**) Quantification and representative immunoblots of striatal, hippocampal and cortical lysates of lamin B1 and (**b, d**) lamin B2 at different stages of the disease progression (W: weeks; I-IV: Vonsattel grades) in (**a, b**) R6/1, (**c, d**) human brain, and their corresponding controls (wild-type (WT) littermates and non-affected individuals (CTL)). Tubulin was used as loading control. Data is expressed as a percentage of controls. Each point corresponds to the value from an individual sample. Bars represent the mean ± S.E.M. Two-tailed unpaired Student’s t test. *p < 0.05, **p < 0.01 and ***p < 0.001 as compared with WT mice or CTL

### Lamin B1 increased levels are mainly localized in neurons

To address the cell-type specificity of the lamin B1 increase we performed NeuN and lamin B1 co-immunostaining (with or without GFAP), in brain sections obtained from 30-week-old R6/1 mice and from HD patients at different stages of the disease. Analysis of confocal z-stacks images from the striatum of R6/1 mice showed a strong increase in lamin B1 signal specifically in NeuN positive nuclei (Fig. 2a). In addition, neuronal nuclei showed clear nuclear lamina invaginations and lamin B1 delocalization within the nucleoplasm as compared to the nuclear periphery pattern observed in wild-type mice striatal neurons. In line with that, neurons from the putamen of HD patients displayed lamin B1 signal delocalization within the nucleus and nuclear invaginations, while nuclei of glial cells were not altered (Fig. 2b). In the hippocampus of R6/1 mice, the dentate gyrus (DG) and the cornu ammonis 1 (CA1) regions presented the highest increase in lamin B1 signal (Fig. 2c). Moreover, CA1 neuronal nuclei displayed morphological alterations and lamin B1 protein delocalization from the nuclear periphery. Interestingly, at early disease stages (12 weeks), these changes appeared to be restricted to the CA1 region (Supplementary Fig. 1d). In addition, altered lamin B1 localization and nuclear morphology was also detected in neurons without mhtt inclusions (Fig. 2d). Overall, these results indicated a cell-type specific increase in lamin B1 levels in response to mutant huntingtin in co-occurrence with lamin B1 delocalization and nuclear lamina morphological alterations independent of the presence of mutant huntingtin aggregates.

**Figure 2.**
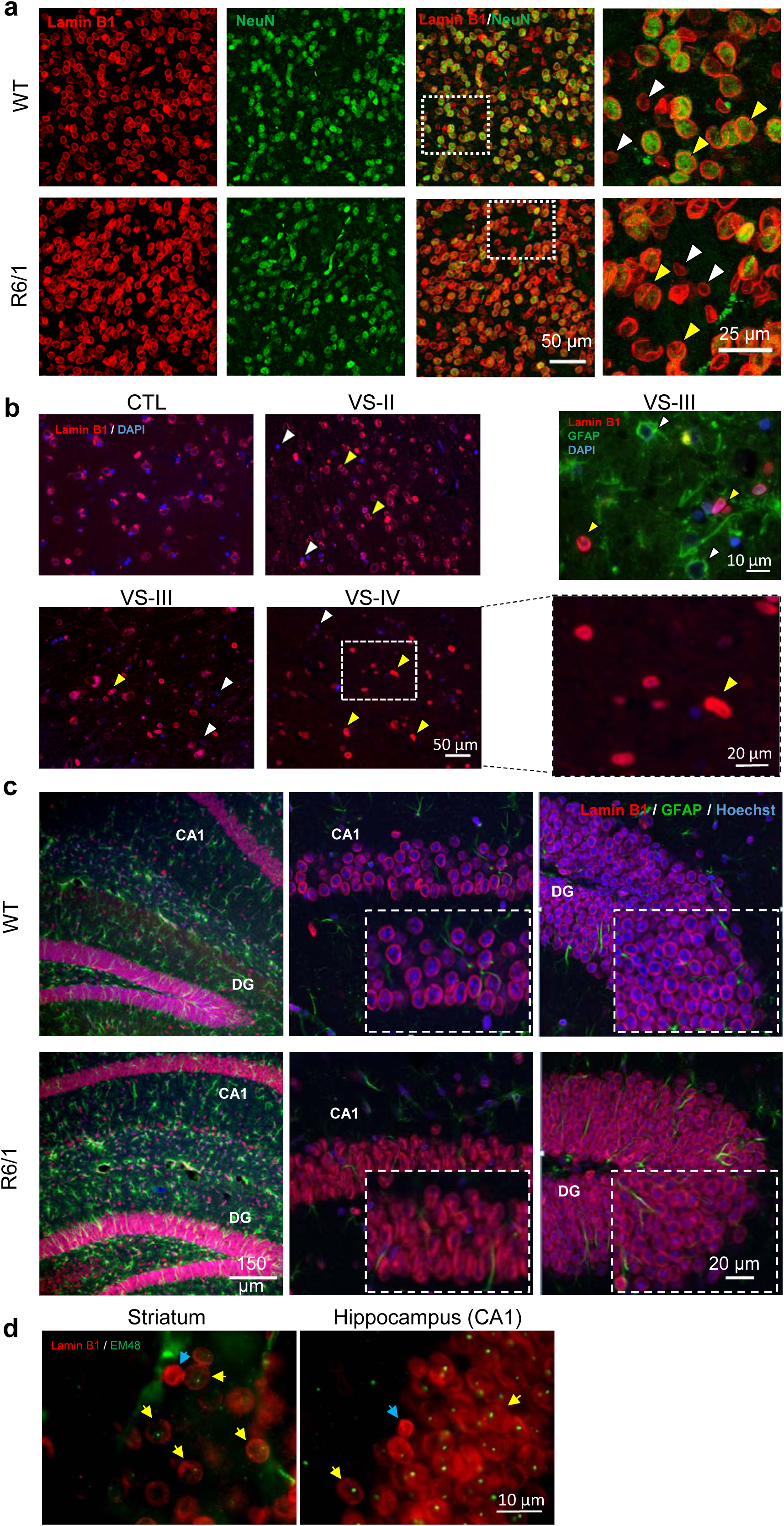
Lamin B1 protein levels are increased mainly in neuronal nuclei. Representative images showing the distribution of lamin B1 in (**a**) The striatum and (**c**) Hippocampus of WT and R6/1 mice at 30 weeks of age and in (**b**) The putamen of HD patients at different stages of the disease (VS: Vonsattel grade). (**a**) Antibody against lamin B1 (red) was combined with anti-NeuN antibody (green) to label neurons nuclei. Yellow arrowheads show co-localization between lamin B1 and NeuN and white arrowheads show lamin B1 positive nuclei not positive for NeuN. Images at high magnification (right) show changes in the nuclear morphology, including nuclear elongation and nuclear envelope invaginations; and in lamin B1 nuclear pattern in the R6/1 mice, presenting appreciable levels of lamin B1 within the nucleoplasm. (**b**) Antibody against lamin B1 (red) was combined with DAPI-Fluoromount G (blue) to label nuclei. Images at low (left and right-up) and high (right-down) magnification show MSNs (yellow arrowheads) and glial cells (white arrowheads). Images show altered nuclear morphology in neurons including nuclear elongation and nuclear envelope invaginations. (**c**) Antibody against lamin B1 (red) was combined with anti-GFAP antibody (green) to labell astrocytes and Hoechst 33258 (blue) to label nuclei. (**d**) Representative images showing lamin B1-positive nuclei (red) and mhtt aggregates labeled with the EM48 antibody (green) in the striatum and hippocampus (CA1) of 30-week-old R6/1 mice. Yellow arrows show nuclei with mhtt aggregates and no changes in lamin B1 levels and nuclear morphology, and blue arrows show nuclei without aggregates and changes in lamin B1 levels and nuclear morphology.

### Cell-type specific nuclear morphology alterations correlate with increased lamin B1 levels

To determine whether lamin B1 protein levels and nuclear morphology are altered in specific cell populations, fluorescence-activated nuclear suspension imaging (FANSI) method was developed by combining nuclear isolation from brain tissue with immunostaining^19^ and the recently developed imaging flow cytometry^20^, allowing the acquisition of individual nuclei images. By combining antibodies against Ctip2 and NeuN we discerned between MSNs (Ctip2^+^/NeuN^+^), which represent around 90% of total striatal neurons^21^, and glial cells (Ctip2^−^ /NeuN^−^; Supplementary Fig. 2a). We detected an increase in lamin B1 levels and altered nuclear morphology in 30-week-old R6/1 MSNs (Fig. 3a), but not in glial cells (Supplementary Fig. 3a). Nuclear area and total number of nuclei were not altered in comparison to wild-type mice, in agreement with the lack of neuronal death observed in R6/1 mouse brain^22^. In the putamen of HD patients, Ctip2^+^/NeuN^+^ nuclei did not display significant differences in lamin B1 levels or nuclear morphology buttheir area was reduced, in Vonsattel grade III-IV HD patients (Fig. 3b). As expected, the number of MSNs nuclei was reduced in the putamen of HD patients in comparison to control individuals (Fig. 3b). In addition, accordingly to results obtained in R6/1 mice striatum, no alterations were detected in striatal glial cells from HD patients in comparison to control individuals (Supplementary Fig. 3b). To distinguish between DG and CA1 mouse hippocampal neuronal nuclei, we used antibodies against Ctip2 and Prox1 (Supplementary Figure 2b). We detected increased lamin B1 levels only in CA1 neuronal nuclei (Ctip2^+^/ Prox1^−^) in concomitance with morphological alterations, while neuronal nuclei from DG (Ctip2^+^/ Prox1^+^) were relatively spared. Furthermore, no alterations were found in the total number of nuclei or in nuclear area (Fig. 3c).

**Figure 3.**
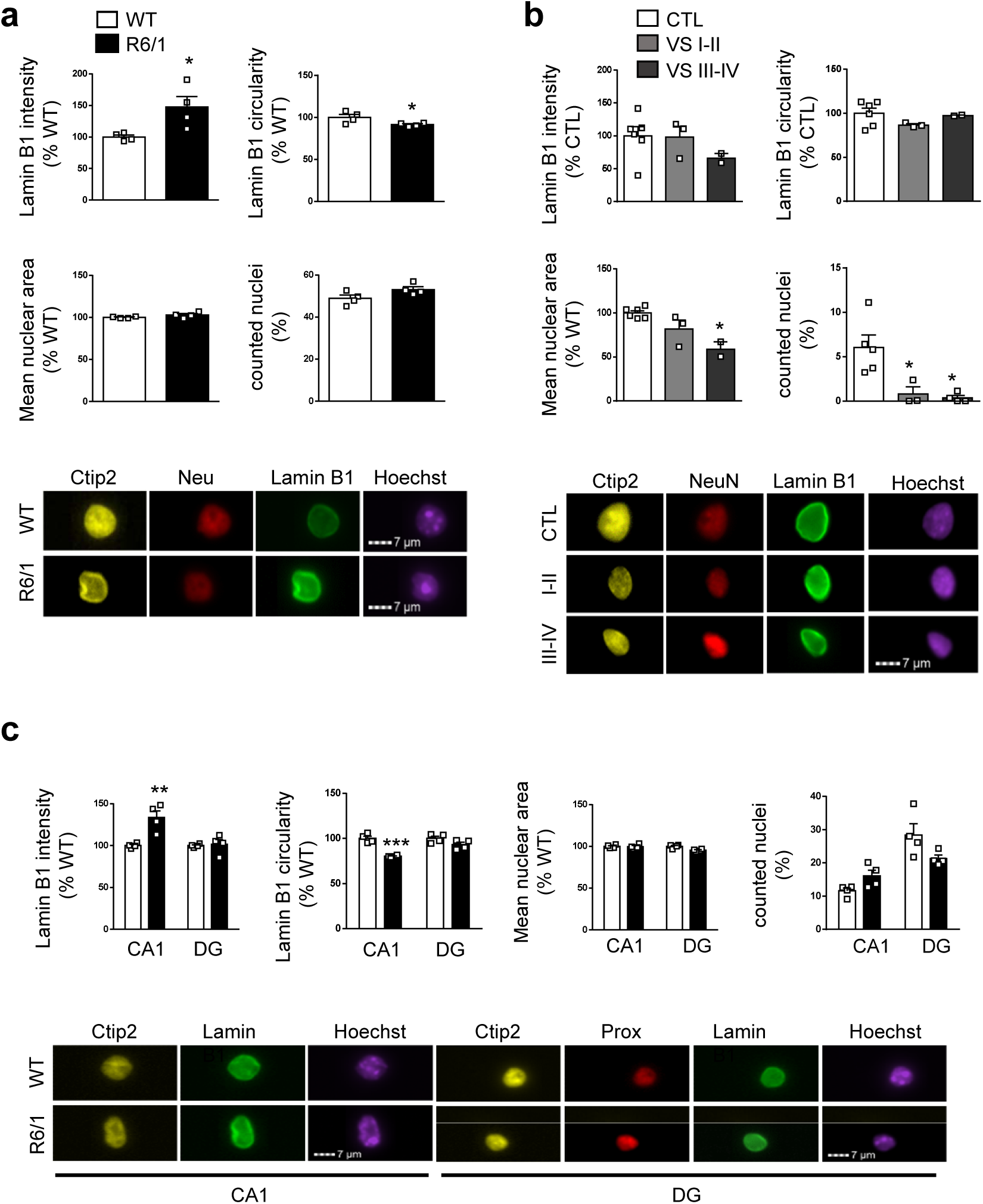
Increase in lamin B1 levels and altered nuclear morphology occur in striatal MSNs and hippocampal CA-1 neurons from R6/1 mice. (**a**) Graphs show the quantification of different parameters in striatal MSNs (Ctip2+/NeuN+) nuclei from 30-week-old wild-type (WT) and R6/1 mice. Representative images are shown. (**b**) Graphs show the quantification of different parameters in striatal MSNs (Ctip2+/NeuN+) nuclei from HD patients at different stages of the disease (Vonsattel grades, VSI-II and III-IV). Representative images are shown. (**c**) Graphs show the quantification of different parameters in hippocampal CA1 (Ctip2+/Prox1-) and DG (Ctip2+/Prox1+) neurons nuclei from wild-type (WT) and R6/1 mice. Representative images are shown. (**a-d**) Each point corresponds to the value from an individual sample. Bars represent the mean ± S.E.M. Two-tailed unpaired Student’s t test. *p < 0.05 as compared with corresponding controls.

### Increased lamin B1 levels correlates with alterations in nuclear permeability in MSNs and CA1 hippocampal neurons from R6/1 mice

Our previous results showed that striatal MSNs and CA1 hippocampal neurons nuclei are preferentially affected by altered lamin B1 levels. Since alterations in nuclear architecture lead to changes in nuclear permeability^23^ we predicted that increased lamin B1 levels could result in nuclear transport abnormalities. To confirm this hypothesis, we performed FRAP experiments in isolated wild-type and R6/1 mice neuronal nuclei by using 20 kDa FITC-dextran (Fig. 4a and Supplementary Fig. 4a). We detected that half-time of recovery (t1/2) after dextran penetration within R6/1 MSNs and CA1 hippocampal, but not DG, nuclei was slower than in wild-type mice (Fig. 4b-d) although bleaching percentage and maximum fluorescence recovered (plateau) did not differ between genotypes (Supplementary Fig. 4b, c). Altogether, our results show an alteration in the entrance of dextran into neuronal nuclei containing greater lamin B1 protein levels, indicating that nucleocytoplasmic passive transport abnormalities are linked to lamin B1 alterations in a cell-type dependent manner.

**Figure 4.**
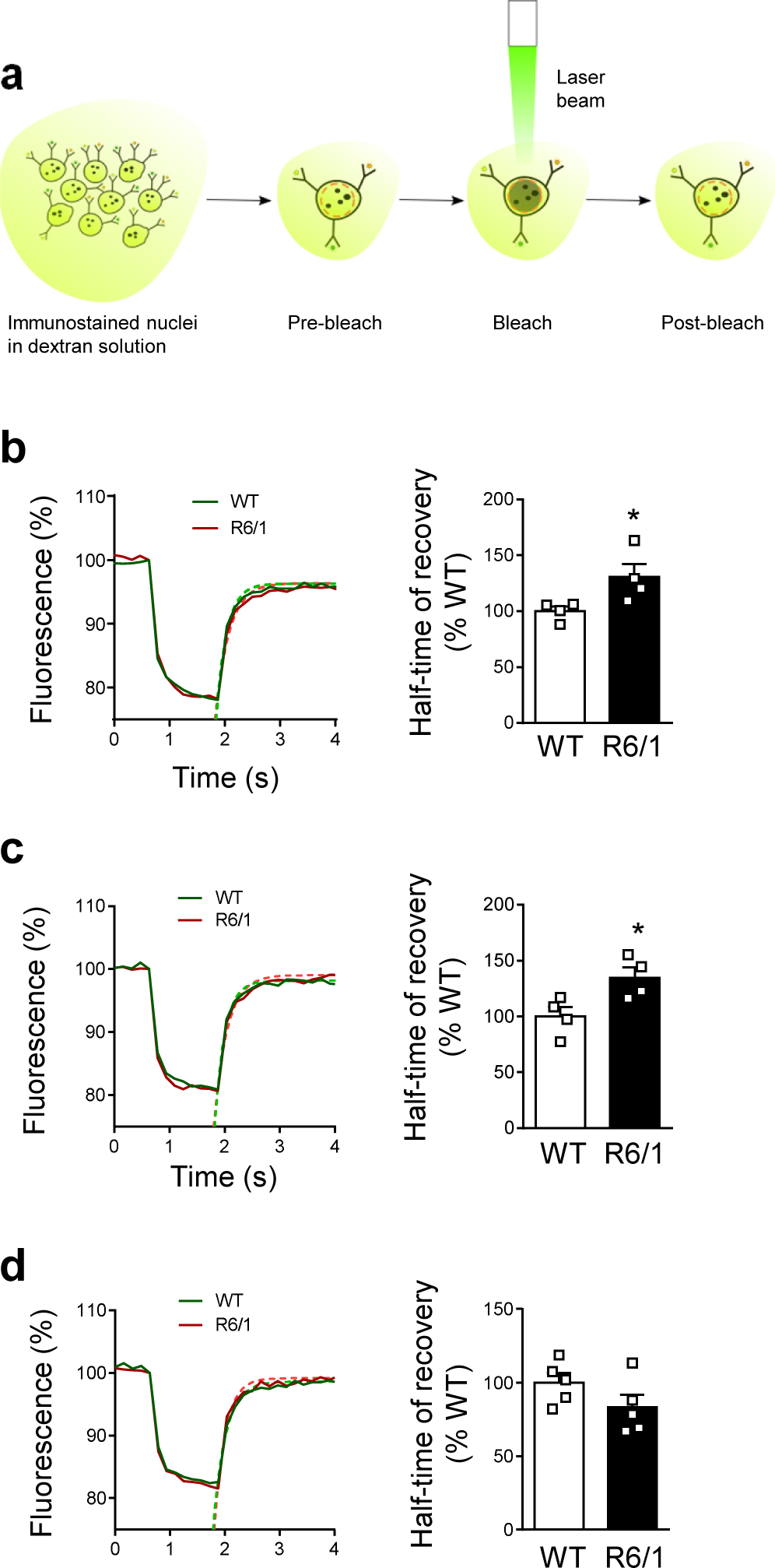
Altered nuclear permeability in striatal MSNs and hippocampal CA1 neurons nuclei from R6/1 mice. (**a**) Scheme showing the experimental approach followed to measure nuclear permeability. (**b**) Nuclear permeability in striatal MSNs nuclei from 30-week-old wild-type (WT) and R6/1 mice. (**c**) Nuclear permeability in hippocampal CA1 neurons nuclei from 30-week-old wild-type (WT) and R6/1 mice. (**d**) Nuclear permeability in hippocampal DG neurons nuclei from 30-week-old wild-type (WT) and R6/1 mice. (**b-d**) Each point corresponds to the value from an individual sample. Bars represent the mean ± S.E.M. Two-tailed unpaired Student’s t test. *p < 0.05 as compared with WT mice.

### Lamin B1 chromatin binding is impaired in R6/1 mice hippocampus

Lamin B1 is classically associated to large heterochromatin domains called lamin associated domains (LADs), characterized by low gene expression levels^24^. Recently, however, LADs have been linked to actively-transcribed euchromatic regions^25^. Therefore, we set out to study whether increased lamin B1 levels could alter their chromatin binding landscape. For that, we generated lamin B1 chromatin immunoprecipitation and sequencing (ChIP-seq) data in 30-week-old wild-type and R6/1 mice hippocampus (age and region showing the highest increase in lamin B1 levels). We immunoprecipitated lamin B1 and verified that both heterochromatin and euchromatin fractions were efficiently sonicated (Supplementary Fig. 5a, b). This indicated that all lamin B1 bound regions should be captured in our ChIP-seq experiments. We ran the EED peak calling tool^26^ and identified 145 and 166 LADs in wild-type and R6/1 mice hippocampus, respectively (Fig. 5a, b). These numbers were consistently found in triplicates of each experiment (Supplementary Fig. 5c, d), and the regions overlapped with LADs previously identified in neural progenitor cells using DamID^27^ (Supplementary Fig. 5e). Our LADs were depleted of H3K9ac and CTCF and highly enriched in H3K9me3, which are major features of canonical gene-silencing LADs (Supplementary Fig. 5f). We observed a lower average size and lamin B1 binding in R6/1 mice specific LADs (Fig. 5a, b). Interestingly, while most of the regions identified by EDD were common between wild-type and R6/1 mice, a small subset of them were specifically identified in one of the genotypes (Fig. 5C). We found regions specific from R6/1 mice, but they were equally enriched in lamin B1 in both genotypes, suggesting an artifactual origin from the EDD tool. For wild-type specific regions, however, we observed a clear reduction of lamin B1 binding in R6/1 mice (Fig. 5c). Genes located within common LADs between wild-type and R6/1 mice were enriched in terms mostly related to ‘olfactory sensory perception’ and ‘keratinization’ (Fig. 5d). However, genes within wild-type exclusively identified regions showed a strong differential functional signature, being mostly enriched in genes related to ‘nucleosome assembly’ (Fig. 5d). In line with this, nuclear fractionation clearly showed a reduction in the proportion of lamin B1 bound to chromatin, with a parallel accumulation of lamin B1 protein within the nucleoplasm (Fig. 5e). Altogether these results suggest that alterations in lamin B1 protein levels and localization in R6/1 mice hippocampus lead to changes in the genome-wide map of LADs, which ultimately could affect the expression and accessibility of certain genes.

**Figure 5.**
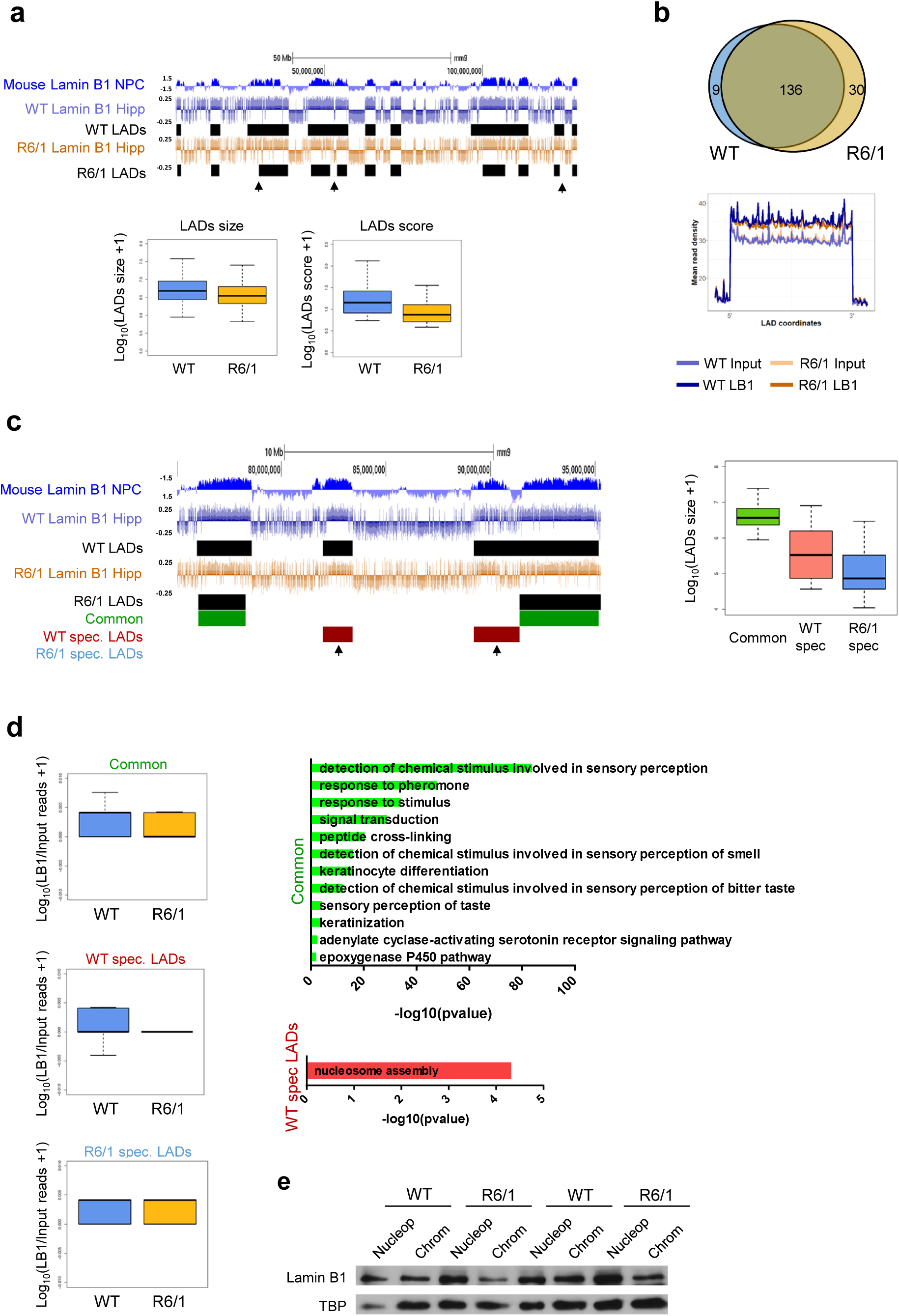
Lamin B1 chromatin binding in wild-type and R6/1 mice hippocampus. UCSC genome browser capture of lamin B1 ChIP-seq signal (log(LB1/Input)) and LADs discovered in wild-type (WT) and R6/1 mice in combination with NPC lamin B1 DamID (left). Boxplots of LADs size (log_10_(LADs size + 1)) and of LADs score (log_10_(LADs score + 1)) obtained by EDD (right). Hipp, hippocampus. (**b**) Venn diagram of overlapping LADs between wild-type (WT) and R6/1 mice hippocampus (top). Metaprofile of lamin B1 and Input datasets mean read density within common LADs for WT and R6/1 mice (bottom). (**c**) UCSC genome browser capture of lamin B1 ChIP-seq signal (log(LB1/Input)) and common, wild-type (WT) specific (spec) and R6/1 specific LADs identified by EDD (left). Boxplot of average size (log_10_(LADs size + 1)) for common, WT specific and R6/1 specific LADs (right). (**d**) Lamin B1 enrichment in common, wild-type (WT) specific (spec) and R6/1 specific LADs (log_10_(LB1/Input reads +1) in hippocampus (Hipp; left). Bargraphs of significant (Benjamini’s adjusted pvalue < 0.05) Biological Processes terms from DAVID for genes within common and WT specific LADs (right). Bars represents the –log_10_ (Benjamini’s adj pvalue). (**e**) Representaive immunoblot showing lamin B1 levels in the nucleoplasm (Nucleop) and chromatin (Chrom) in the hippocampus of 30-week-old wild-type (WT) and R6/1 mice. TBP, TATA binding protein.

### Chromatin accessibility, gene transcription and LADs organization in R6/1 mice hippocampus

To study the impact of lamin B1 chromatin binding alterations in chromatin state and gene expression, we analysed chromatin accessibility and gene expression levels by generating assay for transposase-accessible chromatin and parallel sequencing (ATAC-seq; Fig. 6a and Supplementary Fig. 6a) and RNA-seq data using 30-week-old R6/1 mice hippocampus. We identified a similar number of ATAC-seq peaks in wild-type and R6/1 mice hippocampus (260284 ± 4559 and 254672 ± 9030, respectively) by using three independent biological replicates, indicating no massive changes in chromatin accessibility between genotypes. Differential peak accessibility analysis identified 1304 and 803 regions with gained or lost accessibility, respectively, in R6/1 mice hippocampus (adjusted pvalue< 0.05). These regions were predominantly distal regulatory elements localized at intronic and intergenic regions (Supplementary Fig. 6b). Motif analysis identified EGR1/2 and NEUROD2 as centrally enriched transcription factors in each set of differential accessible peaks (Supplementary Fig. 6c). To gain insight into functional relevance of these changes, differential accessible regions were annotated to the closest transcription start site (TSS). Genes showing a loss of accessibility in R6/1 mice displayed a clear neuronal signature, with enriched terms such as ‘positive regulation of synapse’ or ‘chemical synaptic transmission’, while genes associated with an increase in chromatin accessibility were mostly associated to developmental and transcriptional related terms (Fig. 6b). The transcriptome of both genotypes (using nine replicates per condition) clearly differed (as shown by PCA in Supplementary Fig. 6d). We found 2145 genes up-regulated and 2280 genes down-regulated genes (adjpvalue < 0.001) in R6/1 against wild-type mice hippocampus. Gene ontology analysis was in agreement with the one found in our ATAC-seq data, with a predominance of neuronal-related and transcriptional-related terms for down-and up-regulated genes, respectively (Supplementary Fig. 6e). In addition, we found a substantial overlap with previously identified sets of altered expressed genes in other HD mouse models (Supplementary Fig. 6f)^28,29^. As expected, we found that only genes showing the highest transcriptional dysregulation (adjusted p value <0.001, |fold change| >2) displayed significant changes in chromatin accessibility at their TSS as compared with genes only filtered according to their adjusted p value (Fig. 6c and Supplementary Fig. 6g). However, when analysing transcriptional changes associated with identified differential accessible regions, a clear correlation was observed (Fig. 6d), suggesting that distal regulatory elements accessibility better account for transcriptional dysregulation in R6/1 mice hippocampus.

**Figure 6.**
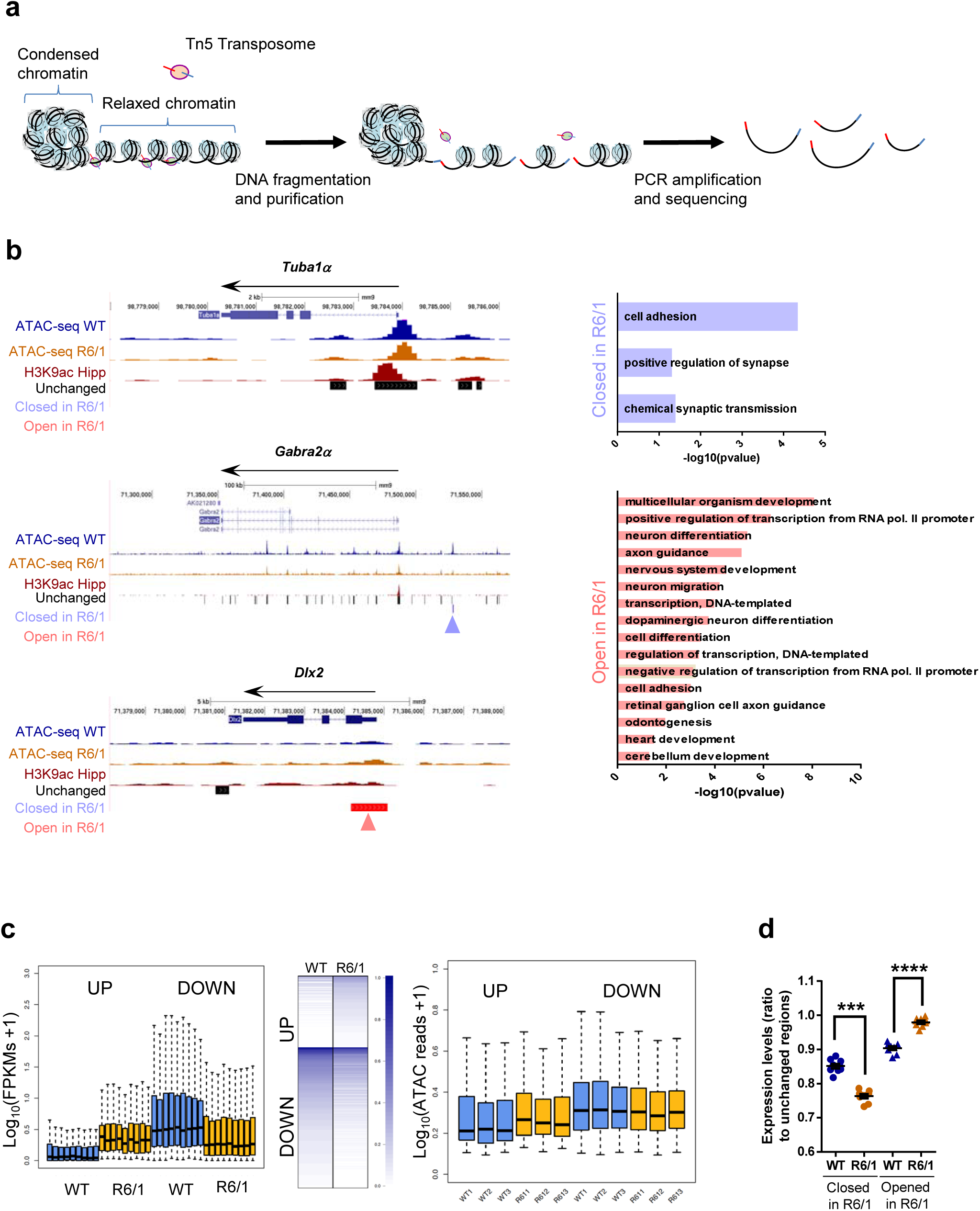
Hippocampal chromatin accessibility and gene expression analysis in R6/1 mice hippocampus. (**a**) Scheme showing major steps of ATAC-seq technique. (**b**) UCSC genome browser capture of wild-type (WT) and R6/1 mice hippocampus ATAC-seq data, hippocampal H3K9ac and unchanged, closed in R6/1 and open in R6/1 accessible detected regions (right) in *Tuba1α, Gabra2α* and *Dlx2* gene locus. Arrows in blue (closed in R6/1) and red (open in R6/1) indicate differential accessible regions. Bargraphs of significant (Benjamini’s adjusted pvalue < 0.05) Biological Processes terms from DAVID for genes associated with decreased (right top) or increased (right bottom) chromatin accessibility regions. Bars represents the –log_10_(Benjamini’s adj pvalue). (**c**) Boxplots showing average gene expression (log_10_(FPKMs +1) for genes up or down-regulated (adjpvalue<0.001, |FC|>2, N=9) in R6/1 mice (left). Heatmap showing expression profile of genes up-or down-regulated (adjpvalue<0.001, |FC|>2, N=9) in R6/1 mice (mid). Boxplots showing average TSS chromatin accessibility (log_10_(ATAC reads +1), N=3) for genes up or down-regulated (adjpvalue<0.001, |FC|>2, N=9) in R6/1 mice hippocampus (right). (**d**) Boxplots showing average gene expression (closed or open regions FPKMs/ unchanged regions FPKMs) for genes associated with differential accessible regions in R6/1 mice (adjpvalue < 0.05, N=3).

When focusing on LADs reported in both genotypes, as expected, we found a particular enrichment in genes with low transcriptional rate (Fig. 7a). However, when we analysed LADs specifically found in wild-type mice or common between both genotypes (Fig. 5c), minor changes were found either at transcriptional level (Fig. 7b) or in terms of chromatin accessibility (Fig. 7c), indicating that loss of lamin B1 binding in R6/1 mice hippocampal cells does not lead to a global transcriptional induction in these regions. Additionally, we analysed the presence of genes differentially expressed within the set of genes specifically found in wild-type LADs (lost in R6/1 mice, see Fig. 5c) or in wild-type and R6/1 mice common LADs (Fig. 7d). We found that genes differentially expressed between wild-type and R6/1 mice were more enriched in wild-type specific LADs than in wild-type and R6/1 common LADs (up-regulated: 119/1242 in wild-type specific LADs vs 142/3654 in common LADs; down-regulated: 110/1242 in wild-type specific LADs vs 198/3654 in common LADs), indicating that loss of lamin B1 chromatin binding in R6/1 mice hippocampal cells could lead to chromatin reorganization affecting genes within these regions. However, when focusing on highly dysregulated genes (|fold change| >2), this enrichment was only observed for down-regulated genes (down-regulated: 10/1242 in wild-type specific LADs vs 21/3654 in common LADs; up-regulated: 3/1242 in wild-type specific LADs vs 9/3654 in common LADs). Furthermore, we analysed whether regions with differential accessibility between wild-type and R6/1 mice displayed changes in lamin B1 chromatin binding. Not surprisingly, ATAC-seq regions were generally depleted of lamin B1 occupancy (Fig. 7e, left panel). However, when comparing regions showing increased or decreased chromatin accessibility in R6/1 mice to unchanged regions, a highest occupancy of lamin B1 was observed in the first, especially in regions with decreased accessibility (Fig. 7e, right panel). Interestingly, a significant decrease in lamin B1 occupancy was observed in regions with increased accessibility in R6/1 mice, suggesting that lamin B1 chromatin binding impairment could indeed lead to localized increase in chromatin accessibility in distal regulatory elements in R6/1 mice hippocampus. Overall, our results suggest that while no massive changes at transcriptional or chromatin accessibility levels are associated to the loss of lamin B1 heterochromatin binding, a small subset of differentially expressed genes could be affected. Moreover, distal regulatory elements, and more particularly those found in regions with gained accessibility in R6/1 mice, appear to be the more sensitive to these lamin B1 alterations.

**Figure 7.**
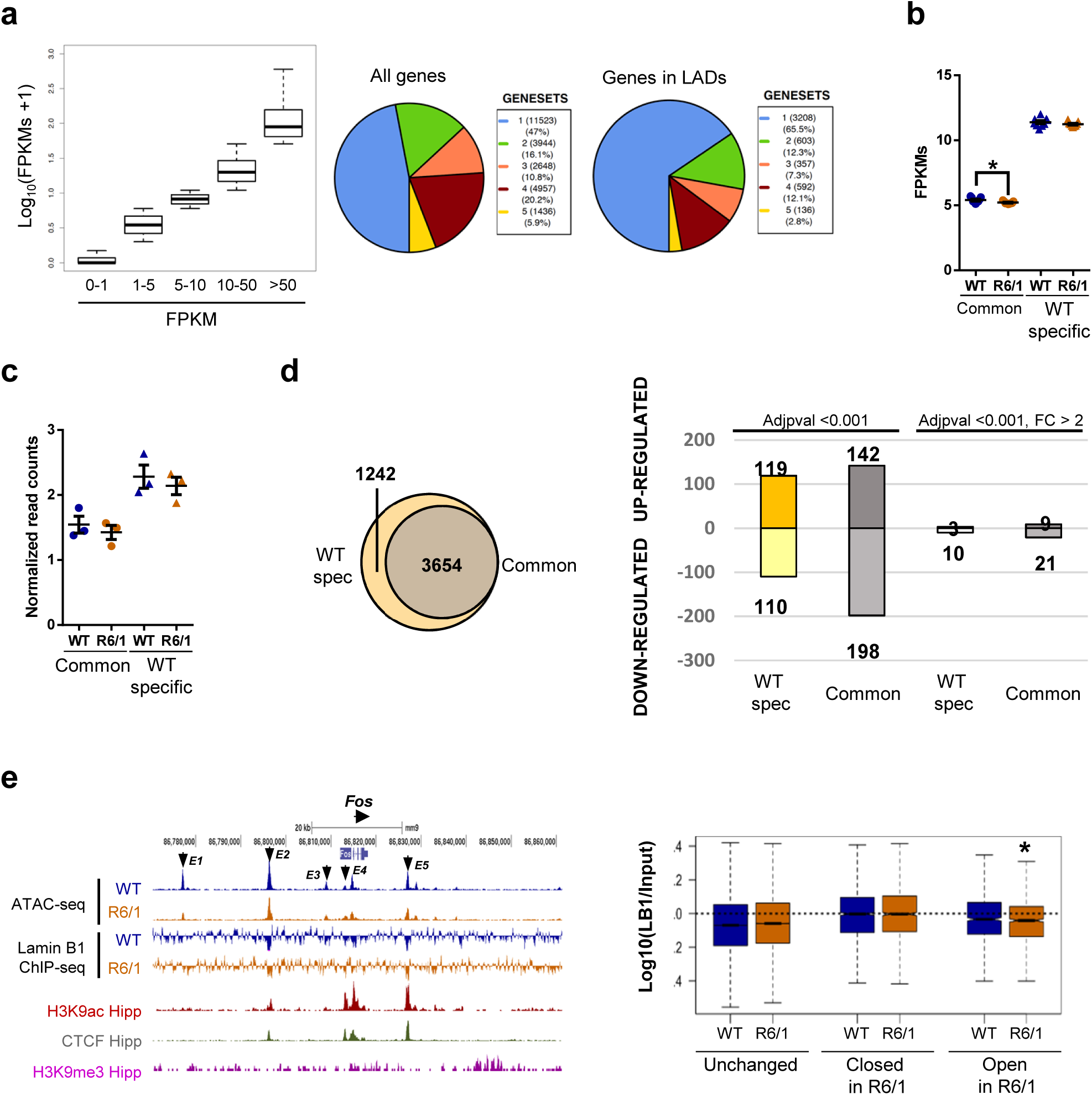
Chromatin accessibility, gene expression and lamin B1 chromatin binding interconnexion. (**a**) Boxplots of average expression (log_10_(FPKMs +1)) for genes sublist 1-5 (lowest to highest expression) for wild-type (WT) mice (left). Pie charts of gene distribution among generated sublists (1-5) for all genes (middle) and genes in LADs (right) in WT mice. Average gene expression (FPKMs) of genes found in common and wild-type (WT) specific LADs for WT and R6/1 mice. Mann-Whitney’s non-parametric t-test was used for statistical analysis; * p<0.05 as compared to WT mice. (**c**) Average chromatin accessibility (normalized read counts) of genes found in common and wild-type (WT) specific LADs for WT and R6/1 mice. Mann-Whitney’s non-parametric t-test was used for statistical analysis. (**d**) Venn diagram showing the total number of genes found in common and wild-type (WT) specific LADs regions(left). Bargraph showing the number of up-and down-regulated genes found in common and WT specific LADs regions filtering only according to adjusted pvalue (adj pvalue <0.001) or additionally with fold change (adjusted pvalue <0.001, |FC| > 2). (**e**) UCSC genome browser capture of wild-type (WT) and R6/1 mice hippocampus ATAC-seq data, lamin B1 ChIP-seq (log(Lb1 ChIP/Input)) for WT and R6/1 mice hippocampus, hippocampal H3K9ac, CTCF and H3K9me3 for WT mice hippocampus in *Fos* locus (right). Enhancer regions (E1-E5) are indicated with arrows. Boxplot of lamin B1 enrichment (log_10_(Lb1 ChIP/Input)) for regions with unchanged, decreased (closed in R6/1) and increased (opened in R6/1) chromatin accessibility in R6/1 mice hippocampus. Mann-Whitney’s non-parametric t-test was used for statistical analysis; * p<0.05 as compared to WT mice.

### Treatment with betulinic acid prevents cognitive impairment in R6/1 mice

Given the increase in lamin B1 levels and the altered nuclear morphology and function found in R6/1 mice brain neurons, we hypothesized that these disturbances could be possibly contributing to motor and cognitive impairment present in HD. Betulinic acid has been shown to transcriptionally repress *Lmnb1* expression^30^. Thus, we treated R6/1 mice from 8 to 20 weeks of age with 50 mg/kg betulinic acid and analyzed behavioral, biochemical and histopathological changes following the timeline depicted in Fig. 8a. First, we analyzed hippocampal-dependent learning and memory by using novel object location test (NOLT) and novel object recognition test (NORT). As previously described^31^, vehicle-treated R6/1 mice showed impaired hippocampal-dependent learning and memory relative to wild-type mice with a decreased percentage of time exploring the moved (Fig. 8b) or the novel (Fig. 8c) objects. Interestingly, betulinic-acid treated R6/1 mice explored similarly to wild-type mice in the NOLT (Fig. 8b) and NORT (Fig. 8c), suggesting that chronic administration of betulinic acid prevents cognitive dysfunction in R6/1 mice. To assess whether learning of a cortico-striatal motor task was also improved after treatment with betulinic acid, we performed the accelerating rotarod task at 15 weeks of age, when R6/1 mice show a clear difference in the performance compared to wild-type mice^31^. Vehicle-treated R6/1 mice displayed poor performance with a lower latency to fall compared to control mice (Fig. 8d). Importantly, betulinic acid-treated R6/1 mice showed a significant, although partial, improvement of their motor learning abilities.

**Figure 8.**
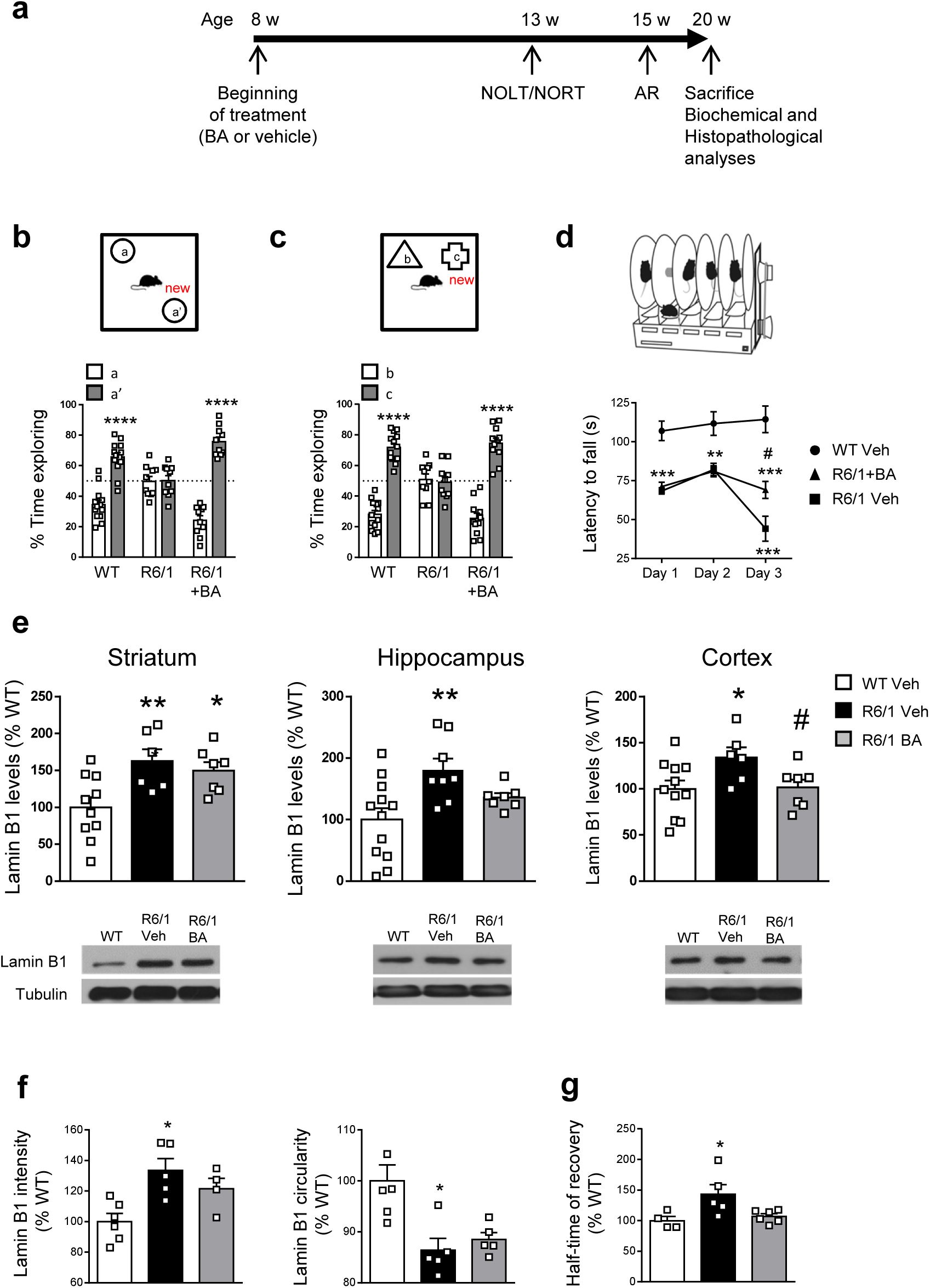
Chronic treatment with betulinic acid improves cognitive function and modulates lamin B1 levels in the hippocampus and cortex of R6/1 mice. (**a**) Timeline of the behavioral, biochemical and histopathological analyses performed in wild-type (WT) and R6/1 mice to assess the effect of betulinic acid (BA) administration. W, weeks; AR, accelerating rotarod. (**b, c**) Graphs show the percentage of time exploring each object with respect to the total exploration time in the (**b**) NOLT and (**c**) NORT, 5 weeks after treatment (Veh, vehicle; BA, betulinic acid; WT, wild-type). ****p < 0.0001 compared to the corresponding old location/object. (**d**) Accelerating rotarod was assessed after 7 weeks of treatment. **p < 0.01 and ***p < 0.001 compared to vehicle-treated wild-type (WT) mice; ^#^p < 0.05 compared to vehicle-treated R6/1 mice. (**e**) Lamin B1 levels in the striatum, hippocampus and cortex after 12 weeks of treatment. *p < 0.05, **p < 0.05 compared to vehicle-treated wild-type (WT) mice and ^#^p < 0.05 compared to vehicle-treated R6/1 mice. Representative immunoblots of lamin B1 and α-tubulin for each treatment group are shown. (**f**) Lamin B1 intensity and nuclear morphology in hippocampal CA1 neurons nuclei from wild-type (WT) and R6/1 mice after 12 weeks of treatment. *p < 0.05 compared to vehicle-treated WT mice. *p < 0.05 as compared to vehicle-treated WT mice. (**g**) Nuclear permeability in hippocampal CA1 neurons nuclei after 12 weeks of treatment. *p < 0.05 compared to vehicle-treated wild-type (WT) mice. *p < 0.05 as compared to vehicle-treated WT mice. In all graphs, bars represent the mean ± S.E.M and each point corresponds to the value from an individual sample Statistical analysis was performed by one-way ANOVA followed by Bonferroni’s post hoc test except in (**d**) where data was analyzed by two-way ANOVA followed by Bonferroni’s *post hoc* test.

Next, we investigated whether chronic betulinic acid treatment affected lamin B1 protein levels in different brain regions. A reduction of lamin B1 protein levels in the cortex and hippocampus, but not in the striatum, was detected in betulinic acid-treated R6/1 mice (Fig. 8e). Analysis using FANSI revealed a normalization of lamin B1 levels in the nuclei of hippocampal CA1 neurons in betulinic acid-treated R6/1 mice (Fig. 8f), accompanied by a partial rescue of nuclear morphology alterations (Fig. 8f). In line with these results, we observed an amelioration of nucleocytoplasmic transport dysfunction, since half-time of recovery after photobleaching of nuclear dextran fluorescence was similar in betulinic-acid treated R6/1 and vehicle-treated wild-type mice hippocampal CA1 neuronal nuclei (Fig. 8g). Importantly, as observed at 30 weeks of age, lamin B1 levels and nuclear morphology or permeability alterations were not detected in hippocampal DG neurons nuclei from 20-week-old vehicle-or betulinic-acid treated R6/1 mice (Supplementary Fig. 7a, b).

Finally, we analyzed whether several hallmarks of the disease were affected by the treatment with betulinic acid. We found that betulinic acid did not prevent the loss in DARPP-32 levels in the striatum of R6/1 mice, whereas it completely prevented the loss in hippocampal PSD-95 levels (Supplementary Fig. 7c, d). Moreover, we found that the number and area of mhtt aggregates in betulinic acid-treated R6/1 mice were similar to those found in the vehicle-treated group, both in the striatum and hippocampus (Supplementary Fig.7e, f), indicating that betulinic acid effects were independent from mhtt aggregation.

## Discussion

Here we show that: (I) lamin B1 protein levels are increased in vulnerable regions of HD brain correlating with altered nuclear morphology; (II) that nucleocytoplasmic transport of small molecules is altered in neurons showing increased lamin B1 levels in R6/1 mouse brain; (III) in R6/1 mice hippocampus (a) lamin B1 alterations lead to partial un-structuring of LADs and (b) changes in chromatin accessibility mostly localize at distal regulatory elements, correlate with transcriptional dysregulation and are partially associated with lamin B1 chromatin binding alterations, and (IV) pharmacologic regulation of lamin B1 levels normalizes nucleocytoplasmic transport alterations and ameliorates multiple behavioral abnormalities of R6/1 mouse.

We found that, among all lamin isoforms, lamin B1 was consistently affected in the brain of both HD patients and mouse models from early stages of the disease. B-type, but not A-type lamins, are essential for brain development^32^. In fact, none of the laminopathies related to mutations in lamin A/C course with neuronal dysfunction which could explain why lamin A/C were unaltered in most R6/1 brain regions. Alterations in B-type lamins have been reported in neurodegenerative disorders and aging. For instance, and in contrast to our results, decreased lamin B levels are found in brains from AD^33^ and PD patients bearing the LRRK2(G2019S) mutation^8^, and in aged primary human fibroblasts^34^ and keratinocytes^35^, being a marker of cellular senescence. Interestingly, autosomal dominant leukodystrophy, a laminopathy caused by the duplication of the *LMNB1* gene, courses with severe central nervous system affectation, whose symptoms recall those of Huntington’s disease^36^. These evidences prompt the idea that increased lamin B1 levels in HD brain may be participating in the pathophysiology of the disease.

The accumulation of lamin B1 in neuronal nuclei from HD brain could be produced by different mechanisms such as increased transcription and/or translation, or decreased degradation. Increased transcription seems improbable since RNA-seq data generated from hippocampus of 30-week old R6/1 mice do not show alterations in lamin B1 RNA levels (present results)^28^. In addition, although we have recently showed increased translation in the striatum of HD mouse models and patients, the proteomic analysis did not reveal lamin B1 as one of the proteins with increased translation^37^. Therefore, we hypothesize that increased lamin B1 protein levels in HD brain could be the result of different altered post-translational mechanisms such as decreased PKCδ^15^ and/or altered autophagy-mediated lamin B1 degradation ^38^ and/or increased stabilization due to over activation of p38MAPK^39^.

Immunohistochemical analysis of lamin B1 in R6/1 mouse brain suggested that the increase occurred specifically in striatal MSNs and in hippocampal CA1 and DG neurons. Our newly developed technique FANSI, confirmed the increase in lamin B1 levels in nuclei of striatal MSNs and showed that in the hippocampus only CA1 neurons were affected. Interestingly, these are the most vulnerable neurons in HD brain and their dysfunction participate in the motor and cognitive phenotype^40,41^. In the putamen of HD patients, FANSI results indicated a significant loss of MSNs, but not of glial cells, as previously described^42^. However, in contrast to what we observed by immunohistochemistry, we did not detect an increase in lamin B1 levels in any of the cellular populations analyzed suggesting that sample processing could be affecting preferentially nuclei with higher lamin B1 levels (i.e. nuclei with higher nuclear morphology alterations). Altogether, these data support the idea of a cell-type dependent increase in lamin B1 levels, specifically in those neurons preferentially affected in HD.

Our results show that increased lamin B1 levels in the R6/1 mice neuronal nuclei correlated with altered nuclear morphology. In agreement with our results: (I) nuclear morphology alterations in autosomal dominant leukodystrophy brain cells have been related to increased lamin B1 levels^17^, and (II) by using lamin B1 as a marker, nuclear envelope abnormalities have been shown in the brain of HD patients and mouse models^10^. Furthermore, decreased levels of lamin B1 also result in nuclear morphology alterations in AD^33^ and PD^8^ neurons suggesting that proper lamin B1 levels are necessary to maintain a correct neuronal nuclei morphology. In addition, nucleocytoplasmic transport was altered in those nuclei with increased lamin B1 levels in accordance with previous literature showing a link between alterations in lamins and nuclear dysfunction. For instance, in Hutchinson-Gilford progeria syndrome, the mutated form of A-type lamin induces nucleocytoplasmic transport dysregulation by inhibiting the nuclear localization of Ubc9 and disrupting the nucleocytoplasmic Ran gradient, necessary for active transport^43^. In addition, cells expressing progerin presented perturbed passive and active transport towards and from the nucleus^44^. Similarly, increased levels of lamin B1 affects nuclear export in HEK293 cells^16^ and reduces nuclear ion channel opening in fibroblasts from autosomal dominant leukodystrophy patients^17^. Furthermore, altered lamin B levels and nucleocytoplasmic transport occur in AD^45^ and PD^46^ although these alterations have never been linked between them. Altered nucleocytoplasmic mRNA transport has been previously reported in other HD models and has been indirectly associated with nucleoporins sequestration by mhtt inclusions^10,11^. In contrast, our immunohistochemical analyses revealed that the presence of nuclear mhtt inclusions did not necessarily correlate with increased lamin B1 levels or alterations in nuclear morphology in R6/1 mice and HD patients’ striatal and hippocampal neurons. These differences may be due to the use of different HD models, in which forms of mhtt aggregation differ^47–49^. Altogether, here we show for the first time in HD, nuclear morphology and nucleocytoplasmic transport abnormalities in a cell-type dependent manner, and, importantly, directly related with lamin B1 protein levels.

In order to study the consequences of lamin B1 alterations in nuclear lamina heterochromatin organization, we generated, for the first time, lamin B1 ChIP-seq data in mouse central nervous system tissue and characterized hippocampal LADs which, as expected, showed high homology with previous identified domains using DamID^27^. Our lamin B1 ChIP-seq data, together with nuclear fractionation experiments, clearly showed a perturbation in nuclear lamina heterochromatin organization and lamin B1 chromatin binding. Interestingly, previous studies demonstrated that lamin B1 over-expression in the central nervous system leads to epigenetic alterations affecting the heterochromatin protein 1 β(HP1β) and methylated histone H3 (K3H9) as well as transcriptional programs mostly linked to glial cells^16^. In agreement with that, regions with gained chromatin accessibility in our ATAC-seq data showed decreased levels of lamin B1, correlative increased expression and were enriched in terms associated with cell division and development, more typically associated to glial than to neuronal cells^50^. While a previous study showed a general gain of chromatin accessibility in HD T-cells^51^, our hippocampal ATAC-seq data showed a bi-directionality in chromatin accessibility changes, with a clear compaction of neuronal-associated regulatory regions and increased chromatin relaxation in developmental related ones. This is in agreement with a recent study demonstrating that HD neuronal and glial cells are affected in opposite ways at transcriptional level^50^. Our R6/1 hippocampal transcriptional data showed a great overlap with previously generated data in additional HD models^29,52^ and, according to a recent study, these common transcriptional signatures present high homology with those found in knockouts for histone acetyltransferases and methyltransferases^28^. Being shown the interplay between lamin B1 protein levels and H3K9me3, highly dependent on the activity of methyltransferases, it can be speculated that nuclear lamina alterations could play a role in a particular subset of the HD transcriptional signature.

Finally, with the purpose of addressing the therapeutic relevance of our findings, we used an FDA approved compound, betulinic acid, that has the potential to normalize lamin B1 protein levels^30^. We show a strong beneficial effect in preventing HD cognitive dysfunction and, for the first time, in normalizing lamin B1 protein levels in the brain *in vivo*. Interestingly, normalization of lamin B1 levels in the hippocampus of R6/1 mice was accompanied by an improvement in nuclear morphology and function of CA1 neurons with a functional recovery of hippocampal memory dependent tasks. In this line, cytotoxicity is reduced in primary cortical neurons expressing mhtt after pharmacological restoration of nucleocytoplasmic transport^11^. Our results strengthen the idea of a relationship between increased lamin B1 levels and alterations in nuclear morphology and function in HD, as previously suggested in autosomal dominant leukodystrophy^17^, which ultimately contribute to the HD phenotype. In contrast, lamin B1 levels were not normalized in the striatum after betulinic acid treatment although amelioration of motor learning dysfunction was observed in R6/1 mice. Since cortical pyramidal neurons project to the striatum and betulinic acid normalized lamin B1 levels in the cortex, we speculate that improvement of cortical neurons function could have beneficial effects on MSNs, reflected by the prevention in the decrease of DARPP-32 protein levels, a hallmark of HD ^53^, and consequently motor performance is partially improved. In addition, betulinic acid has anti-inflammatory effects ^54^ which may contribute to the improvement of striatal MSNs. Therefore, our results open a new therapeutic window not only for HD, but also for autosomal dominant leukodystrophy, for which no effective treatment is available yet^6^.

Altogether, our findings evidenced a relationship between increased lamin B1 levels and nuclear morphological and functional alterations in HD brain neurons, which may contribute to the pathophysiology of the disease and could have promising applications at the therapeutic level.

## Methods

### HD mouse model

Male R6/1 transgenic mice (B6CBA background) expressing the exon-1 of mhtt with 145 CAG repeats and their wild-type littermate controls were used for this study. Mouse genotyping and CAG repeat length determination were performed as previously described^18^. All mice were housed together in numerical birth order in groups of mixed genotypes, and male littermates were randomly assigned to experimental groups. Data were recorded for analysis by microchip mouse number. The animals were housed with access to food and water *ad libitum* in a colony room kept at 19 – 22 °C and 40 – 60% humidity, under a 12:12 light/dark cycle. All procedures were carried out in accordance with the National Institutes of Health Guide for the Care and Use of Laboratory Animals, and approved by the local animal care committee of the Universitat de Barcelona, following European (2010/63/UE) and Spanish (RD53/2013) regulations for the care and use of laboratory animals.

### Post-mortem human brain tissue

Frozen samples (putamen, hippocampus and frontal cortex) and brain slices (5 μm thick sections paraffin-embedded mounted in glass slides) from HD patients and control individuals were obtained from the Neurological Tissue Bank of the Biobank-Hospital Clínic-Institut d’Investigacions Biomèdiques August Pi i Sunyer (IDIBAPS; Barcelona, Catalonia, Spain) following the guidelines and approval of the local ethics committee (Hospital Clínic of Barcelona’s Clinical Research Ethics Committee). Details on the sex, age, CAG repeat length, Vonsattel grade and *post-mortem* delay are found in Supplementary Table 1.

### Pharmacological treatment

R6/1 mice were treated (from 8 to 20 weeks of age) with vehicle (90% water, 10% Polysorbate 80) or betulinic acid (50 mg/kg; Sigma-Aldrich #855057) administered by oral gavage, 3 days/week. Wild-type mice were treated with vehicle. Animal weight was recorded each day of treatment. Days in which treatment and tests were coincident, mice were allowed to recover during 1 h before starting a task. Mice were sacrificed by cervical dislocation 1 h after the last dose. Half of the brain was fixed in 4% PFA for immunostaining analysis and the striatum, hippocampus and cortex from the other half were rapidly removed and stored at – 80°C until analysis.

### Behavioral assessment

*Spatial and recognition memory tests*. NOLT and NORT were used to analyze hippocampal-dependent spatial long-term and object recognition memory, respectively, in wild-type and R6/1 mice at 13 weeks of age as previously described ^31^. In each of the tests, the object preference was measured as the time exploring each object x 100/time exploring both objects. The arena and the objects were rigorously cleaned between animal trials to avoid odors. Animals were tracked with SMART Junior software from Panlab (Barcelona, Spain). Days, in which treatment and tests were coincident, the mice were allowed to recover for 1 hour (h) after the treatment before starting any task.

*Accelerating rotarod*. For the assessment of motor learning dependent on the corticostriatal connectivity, we performed the accelerating rotarod test at 15 weeks of age. The protocol was performed as previously described^31^ The final performance was calculated as the mean latency to fall during the 3 last trials of each day. Days, in which treatment and tests were coincident, the mice were allowed to recover for 1 h after the treatment before starting any task.

### Protein extraction and Western blot analyses

Animals were killed at different ages by cervical dislocation. Brains were quickly removed and the striata, hippocampi and cortex were dissected out and homogenized in lysis buffer. Protein extraction and Western blot analyses were performed as previously described^55^. After incubation with primary and the appropriated horseradish peroxidase-conjugated secondary antibodies (Supplementary table 2), membranes were washed with Tris-buffered saline containing 0.1% Tween 20. Immunoreactive bands were finally visualized using the Western Blotting Luminol Reagent (Santa Cruz Biotechnology #sc-2048) and quantified by a computer-assisted densitometer (Gel-Pro Analyzer, version 4, Media Cybernetics).

### Immunofluorescence

Mice perfusion, brain processing and immunostaining were performed as previously described^15^. For human tissue, the first step was dewaxing and rehydrating the tissue by performing a series of 5 min each: xylene (four times), absolute ethanol (three times), alcohol 96% (three times) and distilled water. The antigen retrieval was performed afterwards by boiling the sections in citrate buffer (10 mM sodium citrate, 0.05% Tween 20, pH 6.0) in a microwave for 20 min. After this, the DAKO Autostainer Plus was used for a blocking step during 15 min at room temperature with a commercial wash buffer from DAKO supplemented with 3% normal goat serum, three washes with phosphate buffered saline (PBS) and the incubation with the primary antibody in the DAKO Real TM antibody diluent (Agilent #S202230-2) for 30 min. After incubation with primary antibodies, sections were washed with PBS and incubated overnight with corresponding secondary antibodies (Supplementary Table 2). Finally, sections were mounted with DAPI Fluoromount-G (Thermo Fisher Scientific #00-4959-52). Negative controls were performed for each primary antibody and no signal was detected in this condition.

### Immunofluorescence imaging and analysis

Immunostained tissue sections were examined by using the Olympus BX60 (Olympus, Tokyo, Japan) epifluorescence microscope coupled to an Orca-ER cooled CCD camera (Hamamatsu Photonics, Hamamatsu, Japan) or the Leica TCS SP5 laser scanning confocal microscope (Leica Microsystems Heidelberg GmbH, Manheim, Germany) with Argon and HeNe lasers coupled to a Leica DMI6000 inverted microscope at different magnifications (from 10x to 63x). Confocal images were taken as stacks differed in 0.5 μm in Z axis with a HCX PL APO lambda blue 63x numerical aperture objective and standard pinhole (1 Airy disk) and their reconstruction was performed using Image J software (NIH, Bethesda, USA).

### Immunohistochemistry for mhtt aggregates detection

Coronal sections (30 µm) of the whole brain were obtained as described above. Mhtt aggregates detection was performed as previously described^31^ by using the anti-EM48 antibody. EM48 staining was examined in eight slices per animal separated by 240 µm (covering the entire striatum or CA1 hippocampal region) by using the Computer-Assisted Stereology Toolbox (CAST) software (Olympus Danmark A/S, Ballerup, Denmark). Images were analyzed using CellProfiler Analyst software.

### FANSI

FANSI was the result of combining nuclear purification and immunostaining^19^ with ImageStream imaging flow cytometer technology^20^ (Luminex Corporation). *Nuclear purification and immunostaining*. Frozen tissue was homogenized in low sucrose buffer (LSB; 0.32 M sucrose, 5 mM CaCl_2_, 5 mM Mg(Ac)_2_, 0.1 mM EDTA, 50 mM HEPES pH 8.0, 1 mM DTT, 0.1% Triton X-100) and fixed in 1% formaldehyde for 10 min at room temperature in a rotating wheel. Formaldehyde was quenched with 125 mM glycine incubation during 5 min at room temperature in the rotating wheel. Tissue homogenate was collected by centrifugation, resuspended in LSB and mechanically homogenized. After that, the homogenized solution was layered on the top of a high sucrose buffer (1 M sucrose, 3 mM Mg(Ac)_2_, 10 mM HEPES, pH 8.0, 1 mM DTT) and centrifuged at 4 °C to recover the nuclei. These were then resuspended in PBTB (PBS, 5% BSA, 0.1% Tween-20) containing the nuclear antibodies and 3% NHS, and incubated in a rotating wheel at 4 °C during 30 min. After that, samples were washed twice with PBTB plus 3% NHS and stained with corresponding secondary antibodies in PBTB plus 3% NHS at 4°C during 15 min. Nuclei were then washed, stained with Hoechst 33258 (1:10000, Thermo Fisher Scientific #H3569) and directly processed for Imaging flow cytometry. *Imaging flow cytometry (Imagestream)*. Purified nuclei were resuspended in 100 µl and filtered using cells strainers of 50 µm pore size (Sysmex Partec, Kobe, Japan) and posteriorly sorted and imaged using a 60x objective at a maximum speed of 600 nuclei/s depending on the sample concentration. For each replicate a minimum of 10000 nuclei were recorded. Fluorescent minus one controls were used evaluate the specificity of the defined populations by individually removing primary but not secondary antibodies (Supplementary Fig. 8a-f) Recorded files were processed and analyzed by the IDEAS Software provided by the Imagestream machine’s manufacturers (Luminex, Austin, USA) after cross-channel signal compensation. After selecting individual nuclei (singlets, only focused acquired images were used for posterior analysis (Supplementary Fig. 8g-h). Lamin B1 positive nuclei were selected for posterior analysis (Supplementary Fig. 8i) and screened according to their Ctip2 and Prox1 signal and the specific neuronal nuclear marker NeuN. For hippocampal samples, CA1 nuclei were classified as Ctip2^+^/Prox1^−^ DG ones as Ctip2^+^/Prox1^+56,57^. For striatum and putamen, nuclei were classified as MSNs (NeuN^+^/Ctip2^+^)^58,59^ or glia (NeuN^−^/Ctip2^−^). All the features analyzed (mean intensity, circularity and mean area) were performed using these selected populations.

### FRAP in isolated nuclei

Striatal nuclei were isolated as previously described^60^ and then incubated for 30 min at 4°C with the corresponding primary and secondary antibodies (Supplementary Table 2). Nuclei were maintained in LSB until analysis. To perform conventional single-photon FRAP experiments, nuclei were incubated with a solution of 20 kDa FITC-dextran (0.3mg/ml; Sigma-Aldrich #FD20) and seeded in glass bottomed chambers and covered with a cover slip. Striatal MSN nuclei (Ctip2^+^/NeuN^+^) and hippocampal CA1 (Ctip2^+^/Prox1^−^) and DG (Ctip2^+^/Prox1^+^) nuclei were manually selected. Each FRAP experiment started with 5 pre-bleach image scans, followed by 8 bleach pulses of 156 ms each on a spot with a diameter of 2.5 µm in the center of the nucleus. At the post-bleach period, a series of 100 single section images were collected at 156 ms intervals (Supplementary Figure 9). A total of 113 images were acquired for each nucleus, and an average of 25 nuclei were acquired for each animal. Image size was 256 x 56 pixels and the pixel width was 120 x 60 nm. For imaging, the laser power was attenuated to 3% of the bleach intensity. FRAP experiments were performed on a Leica TCS SP5 laser scanning confocal spectral microscope (Leica Microsystems, Heidelberg, Germany) equipped with Argon laser and Leica DMI6000 inverted microscope. Images were acquired using a 63X, 1.4 NA oil immersion objective lens and 1.5 Airy units as pinhole. Image processing was performed using LAS AF Lite Software (Leica Microsystems, Heidelberg, Germany). For each image, the fluorescence in the bleached region was normalized for the fluorescence of the background and the percentage of the initial fluorescence was calculated for each time point. FRAP recovery curves were represented (Supplementary Fig. 9) following the formula 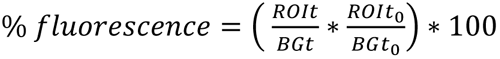 where ROIt is the intensity in the selected ROI at time point t, BGt is the intensity in the background at time point t, ROIt_0_ is the intensity in the selected ROI at time 0, and BGt_0_ is the intensity in the background at time 0. Data were analyzed using Graphpad Prism Software (San Diego, USA).

### Nuclear fractionation

Nuclear fractionation from hippocampus of 30-week old wild-type and R6/1 mice was performed using a Subcellular Protein Fractionation Kit for Tissues (Thermo Fisher Scientific #87790) following manufacturer’s instructions. Chromatin-bound and nuclear soluble fractions were obtained and examined by Western blot as described above.

### RNA-seq

RNA-seq data was generated from 30-week-old wild-type and R6/1 mice hippocampal tissue. RNA was isolated from 9 independent biological replicates for each genotype using the RNeasy Plus Kit (Qiagen #74136) according to the manufacturer’s instructions. Quality assessment was performed using Bioanalyser eukaryiotic total RNA nano series II chip (Agilent #5067-1511) and all samples achieved a RNA integration number (RIN) between 9-10. Libraries were prepared from 9 biological replicates of each condition using the Truseq Stranded mRNA Library Prep Kit (Illumina #20020594) following manufacturer’s instructions and sequenced using the HiSeq-2500 sequencing platform (Illumina, San Diego, USA).

### ChIP-seq

Lamin B1 ChIP-seq was performed as previously described ^61^ by using 30-week-old wild-type and R6/1 mice hippocampal tissue. Briefly, for each biological replicate, hippocampus from 5 mice were pooled together, homogenized in PBS supplemented with proteinase inhibitors (Sigma #13317600) and posteriorly cross-linked with formaldehyde 1% for 15 minutes at room temperature. Cross-linking reaction was stopped by a 5 minutes incubation with 2 M glycine and the cross-linked material was washed 3 times with ice-cold PBS. Cells were lysed using cell lysis buffer (10 mM Hepes pH8, 85mM KCl, 0.5% NP-40) and nuclei were extracted using nuclei extraction buffer (0.5% SDS, 10mM EDTA pH8, 50mM Tris-HCl pH8). Purified nuclear fraction was subjected to sonication using Bioruptor Pico (Diagenode, Belgium) to obtain DNA fragments of 200-500 bp. Sonicated chromatin was incubated overnight at 4°C with anti-rabbit magnetic Dynabeads (Thermo Fisher Scientific #11203D) pre-complexed with 10 µg of rabbit anti-lamin B1 antibody. After 6 washes with RIPA buffer (20 mM Tris-HCl (pH 7.5), 150 mM NaCl, 1 mM Na2EDTA, 1 mM EGTA, 1% NP-40, 1% sodium deoxycholate, 2.5mM sodium pyrophosphate, 1mM b-glycerophosphate, 1mM Na3VO), chromatin was eluted and de-crosslinked by overnight incubation at 65°C, followed by a 30 minutes RNAse (Ambion #AM2271) and 2 hours Proteinase K (Thermo Fisher Scietific #AM2548) treatments. DNA purification was carried out with MinElute PCR Purification KIT (Qiagen #28006) and libraries were prepared using the NEBNext Ultra II DNA Library Prep Kit from Illumina (New England Biolabs #37645) according to the manufacturer’s instructions. DNA size selection was performed after PCR amplification using E-Gel Precast Agarose Electrophoresis System (Invitrogen #A42100). Samples were sequenced single-end using 50 bp reads on the HiSeq-2500 and Hi-Seq 4000 platforms (Illumina, San Diego, USA).

### ATAC-seq

ATAC-seq experiments were performed as previously described^62^, with slight modifications, in 3 independent biological replicates using hippocampal tissue from 25-week-old wild-type and R6/1 mice. Briefly, a frozen mouse hippocampi for each biological replicate was pulverized using a grinder and pestle settle on dry ice, and tissue powder was lysed in LB1 buffer (1M HEPES pH 7.5, 5M NaCl, 0.5M EDTA pH 8.0, 50% glycerol, 10% NP-40, 10% Triton X-100) for nuclear isolation. Approximately 50.000 nuclei were used for the transposition reaction using hyperactive Tn5 transposase (Illumina Cat #FC-121-1030) followed by 13 cycles of PCR amplification. “Nucleosome free” and “mono-nucleosome fragments” were obtained by size-selection of DNA fragments between 170 and 400 bp using SPRIselect beads (Beckman Coulter #B23319) before single-end sequencing to generate 50 bp reads on the Hi-Seq 4000 platform (Illumina, San Diego, USA).

### RNA-seq, ChIP-seq and ATAC-seq data analysis

ChIP-seq samples were mapped against the mm9 mouse genome assembly using Bowtie with the option –m 1 to discard those reads that could not be uniquely mapped to just one region^63^. We ran the EDD tool (parameters: GAP=5 and BIN SIZE=37) to identify LADs on our ChIP-seq samples^26^. Triplicates of each condition were pooled together, once a high degree of similarity in the set of reported LADs and target genes was confirmed among replicates. The UCSC genome browser was used to generate the screenshots of each group of experiments along the manuscript^64^. The RNA-seq samples were mapped against the mm9 mouse genome assembly using TopHat^65^ with the option –g 1 to discard those reads that could not be uniquely mapped in just one region. DESeq2 ^66^ was run over nine replicates of each genotype to quantify the expression of every annotated transcript using the RefSeq catalog of exons and to identify each set of differentially expressed genes. ATAC-seq samples were mapped against the mm9 mouse genome assembly using Bowtie with the option –m 1 to discard those reads that could not be uniquely mapped to just one region, and with the option –X 2000 to define the maximum insert size for paired-end alignment ^63^. Mitochondrial reads were removed from each resulting map and down-sampling was applied to obtain the same number of mapped fragments per sample. Correlation between biological replicates in terms of peaks was assessed to ensure high reproducibility before pooling each set of triplicates. MACS was run with the default parameters but with the shift size adjusted to 100 bp to perform peak calling^67^. The genome distribution of each set of peaks was calculated by counting the number of peaks fitted on each class of region according to RefSeq annotations^68^. Distal region is the region within 2.5 Kbp and 0.5 Kbp upstream of the transcription start site (TSS). Proximal region is the region within 0.5 Kbp upstream of the TSS. UTR, untranslated region; CDS, protein coding sequence; intronic regions, introns; and the rest of the genome, intergenic. Peaks that overlapped with more than one genomic feature were proportionally counted the same number of times. For the generation of metaprofiles, seqminer tool^69^ was used in combination with ggplot2 package from R (https://www.R-project.org/) using an in-house script.

### Differential chromatin accessibility

Integrated analysis of ATAC-seq data was performed using the open Galaxy platform (https://usegalaxy.org/). A list of high confident peaks identified with MACS2 for each genotype was generated by selecting peaks found in at least 2 different replicates. For differential accessible regions identification, edgeR galaxy tool was used with a merge of all high confident peaks identified in both genotypes applyintg the TMM method implemented in edgeR for normalisation and dispersion calculation of the different biological samples. The results were further filtered based on FDR < 0.05. Peaks were annotated to the closest TSS using Homer tool integrated in Galaxy platform.

### Motif analysis

For motif analysis, the Meme-ChIP suite (version 4.12.0) tool^70^ was used in differential enrichment mode together with Hocomoco (version 11 FULL) human and mouse PWMs. As input, 600bp regions surrounding the summit of differential accessible peaks were used for motif discovery (using relaxed regions as control for compacted regions and vice-versa) and only motifs centrally enriched were considered.

### Gene Ontology

For functional enrichments in biological processes (BP) DAVID ^71^ tool was used by providing closest genes ID obtained by Homer when using differentially accessible regions, with subsets of genes identified as differentially expressed or with genes located in LADs identified regions. Terms with Benjamini’s adjusted pvalue <0.05 were selected for bargraph representations.

### Data visualization

UCSC genome browser^64^ was used for genome wide visualization of ChIP-seq and ATAC-seq data.

## Data availability

Raw data and processed information of the ChIP-seq, ATAC-seq and RNA-seq experiments generated in this article were deposited in the National Center for Biotechnology Information Gene Expression Omnibus (NCBI GEO)^72^ repository under the accession number GSE139884. Additionally, public datasets for hippocampal H3K9me3 (GSM2460430)^73^, H3K9ac (GSM2415914)^74^ and CTCF (GSM2228526)^75^ ChIP-seq experiments were retrieved for integrated data analysis.

## Statistics

All the results are expressed as the mean ± SEM. Statistical tests were performed using the Student’s t test for one grouping variable and the one or two-way ANOVA for multi-component variables, followed by Bonferroni’s post hoc test as indicated in the figure legends. A 95 % confidence interval was used and values with a p < 0.05 were considered as statistically significant.

## Acknowledgements

We thank Ana López and Maria Teresa Muñoz for technical assistance; Isabel Crespo from Citomics core facility of the Institut d’Investigacions Biomèdiques August Pi i Sunyer (IDIBAPS) and Maria Calvo from the Advanced Microscopy Unit, Scientific and Technological Centers, University of Barcelona, for their support and advice concerning Cytometry and confocal techniques, respectively; Neurological Tissue Bank of the Biobank-Hospital Clínic-IDIBAPS for providing human brain tissue, and to Ana Saavedra for helpful discussions. Participant labs are supported by: (1) E.P.-N.; Ministerio de Economia y Competitividad, Spain (SAF2016-08573-R) and Fundación Ramón Areces; (2) M.N.; Cancer Research UK Cambridge Institute Core Grant (C9545/A29580); (3) L. DiC.; Ministerio de Economia, Indústria y Competitividad (MEIC; BFU2016-75008-P); (4) S.P.; Instituto de Salud Carlos III (ISCIII; PI18/00283), La Caixa Foundation and Cellex Foundation who provided research facilities and equipment; M.G.-F. was supported by a grant (FI-2016) from Agència de Gestió d’Ajuts Universitaris i de Recerca (AGAUR) and A.P. is a fellow of Sir Henry Wellcome (215912/Z/19/Z).

## Authors contribution

R. A-V., M. G-F. and E. P-N. conceptualized the study and interpreted the results, and E. P.-N. supervised it. R. A-V. and M. G-F. designed and performed most the experiments with the help of J. C.-M., C. C-P. and K. C-V. M. N. conceptualized and contributed to the design of ChIP-, ATAC-and RNA-seq experiments. Y. I. and A. P. contributed with the design and performance of ChIP-, ATAC-and RNA-seq experiments. G. S. and S. S contributed in the analysis of ChIP-, ATAC-and RNA-seq. E. B. analyzed and composed the figures for ChIP-, ATAC-and RNA-seq experiments. L. di C. and S. P. reviewed the content and contributed in the ChIP-, ATAC-and RNA-seq experiments conceptualization and interpretation of the results. R. A-V. and M. G-F., analyzed data and composed the figures. R. A-V., M. G-F. and E. P-N. wrote the manuscript. All the authors critically reviewed the content and approved the final version.

## References

1. de Leeuw, R., Gruenbaum, Y. & Medalia, O. Nuclear Lamins: Thin Filaments with Major Functions. Trends Cell Biol. 28, 34–45 (2018).

2. Hozak, P., Sasseville, A. M. J., Raymond, Y. & Cook, P. R. Lamin proteins form an internal nucleoskeleton as well as a peripheral lamina in human cells. J. Cell Sci. 108, 635–644 (1995).

3. Verstraeten, V., Broers, J., Ramaekers, F. & van Steensel, M. The Nuclear Envelope, a Key Structure in Cellular Integrity and Gene Expression. Curr. Med. Chem. 14, 1231–1248 (2007).

4. Harborth, J., Elbashir, S. M., Bechert, K., Tuschl, T. & Weber, K. Identification of essential genes in cultured mammalian cells using small interfering RNAs. J. Cell Sci. 114, 4557–4565 (2001).

5. Schreiber, K. H. & Kennedy, B. K. When lamins go bad: Nuclear structure and disease. Cell 152, 1365–1375 (2013).

6. Padiath, Q. S. Autosomal Dominant Leukodystrophy: A disease of the nuclear lamina. Front. Cell Dev. Biol. 7, 41 (2019).

7. Hegele, R. A. et al. Sequencing of the reannotated LMNB2 gene reveals novel mutations in patients with acquired partial lipodystrophy. Am. J. Hum. Genet. 79, 383–389 (2006).

8. Liu, G. H. et al. Progressive degeneration of human neural stem cells caused by pathogenic LRRK2. Nature 491, 603–607 (2012).

9. Frost, B. Alzheimer’s disease: An acquired neurodegenerative laminopathy. Nucleus 7, 275–283 (2016).

10. Gasset-Rosa, F. et al. Polyglutamine-Expanded Huntingtin Exacerbates Age-Related Disruption of Nuclear Integrity and Nucleocytoplasmic Transport. Neuron 94, 48–57.e4 (2017).

11. Grima, J. C. et al. Mutant Huntingtin Disrupts the Nuclear Pore Complex. Neuron 94, 93–107.e6 (2017).

12. HDCRG. A novel gene containing a trinucleotide repeat that is expanded and unstable on Huntington’s disease chromosomes. Cell 72, 971–983 (1993).

13. Ross, C. A. & Poirier, M. A. Protein aggregation and neurodegenerative disease. Nat. Med. 10, S10 (2004).

14. Puigdellívol, M., Saavedra, A. & Pérez-Navarro, E. Cognitive dysfunction in Huntington’s disease: mechanisms and therapeutic strategies beyond BDNF. Brain Pathol. 26, 752–771 (2016).

15. Rué, L. et al. Early down-regulation of PKCδ as a pro-survival mechanism in Huntington’s disease. NeuroMolecular Med. 16, 25–37 (2014).

16. Lin, S.-T. & Fu, Y.-H. miR-23 regulation of lamin B1 is crucial for oligodendrocyte development and myelination. Dis. Model. Mech. 2, 178–188 (2009).

17. Ferrera, D. et al. Lamin B1 overexpression increases nuclear rigidity in autosomal dominant leukodystrophy fibroblasts. FASEB J. 28, 3906–3918 (2014).

18. Mangiarini, L. et al. Exon I of the HD gene with an expanded CAG repeat is sufficient to cause a progressive neurological phenotype in transgenic mice. Cell 87, 493–506 (1996).

19. Benito, E. et al. HDAC inhibitor-dependent transcriptome and memory reinstatement in cognitive decline models. J. Clin. Invest. 125, 3572–3584 (2015).

20. Barteneva, N. S. & Vorobjev, I. A. Quantitative Functional Morphology by Imaging Flow Cytometry. Methods Mol Biol 1389, 3–11 (2016).

21. Kemp, J. M. & Powell, T. P. S. The Structure of the Caudate Nucleus of the Cat: Light and Electron Microscopy. Phil. Trans. R. Soc. Lond. B. 262, 383–401 (1971).

22. Francelle, L., Galvan, L. & Brouillet, E. Possible involvement of self-defense mechanisms in the preferential vulnerability of the striatum in huntington’s disease. Front. Cell. Neurosci. 8, 1–13 (2014).

23. Hatch, E. & Hetzer, M. Breaching the nuclear envelope in development and disease. J. Cell Biol. 205, 133–141 (2014).

24. Belmont, A. S., Zhai, Y. & Thilenius, A. Lamin B distribution and association with peripheral chromatin revealed by optical sectioning and electron microscopy tomography. J. Cell Biol. 123, 1671–1685 (1993).

25. Pascual-Reguant, L. et al. Lamin B1 mapping reveals the existence of dynamic and functional euchromatin lamin B1 domains. Nat. Commun. 9, 3420 (2018).

26. Lund, E., Oldenburg, A. R. & Collas, P. Enriched domain detector: A program for detection of wide genomic enrichment domains robust against local variations. Nucleic Acids Res. 42, e92 (2014).

27. Peric-Hupkes, D. et al. Molecular Maps of the Reorganization of Genome-Nuclear Lamina Interactions during Differentiation. Mol. Cell 38, 603–613 (2010).

28. Hervás-Corpión, I. et al. Early alteration of epigenetic-related transcription in Huntington’s disease mouse models. Sci. Rep. 8, 1–14 (2018).

29. Langfelder, P. et al. Integrated genomics and proteomics define huntingtin CAG length-dependent networks in mice. Nat. Neurosci. 19, 623–633 (2016).

30. Li, L. et al. Lamin B1 is a novel therapeutic target of betulinic acid in pancreatic cancer. Clin. Cancer Res. 19, 4651–4661 (2013).

31. Garcia-Forn, M. et al. Pharmacogenetic modulation of STEP improves motor and cognitive function in a mouse model of Huntington’s disease. Neurobiol. Dis. 120, 88–97 (2018).

32. Kim, Y. et al. Mouse B-Type Lamins Are Required for Proper Organogenesis But Not by Embryonic Stem Cells. Science (80-.). 334, 1706–1710 (2011).

33. Frost, B., Bardai, F. H. & Feany, M. B. Lamin Dysfunction Mediates Neurodegeneration in Tauopathies. Curr. Biol. 26, 129–136 (2016).

34. Freund, A., Laberge, R. M., Demaria, M. & Campisi, J. Lamin B1 loss is a senescence-associated biomarker. Mol. Biol. Cell 23, 2066–2075 (2012).

35. Dreesen, O. et al. Lamin B1 fluctuations have differential effects on cellular proliferation and senescence. J. Cell Biol. 200, 605–617 (2013).

36. Padiath, Q. S. et al. Lamin B1 duplications cause autosomal dominant leukodystrophy. Nat. Genet. 38, 1114–1123 (2006).

37. Creus-Muncunill, J. et al. Increased translation as a novel pathogenic mechanism in Huntington’s disease. Brain 142, 3158–3175 (2019).

38. Dou, Z. et al. Autophagy mediates degradation of nuclear lamina. Nature 527, 105–109 (2015).

39. Barascu, A. et al. Oxidative stress induces an ATM-independent senescence pathway through p38 MAPK-mediated lamin B1 accumulation. EMBO J. 31, 1080–1094 (2012).

40. Vonsattel, J. G. & DiFiglia, M. Huntington Disease. J. Neuropathol. Exp. Neurol. 57, 369–384 (1998).

41. Murphy, K. P. et al. Abnormal synaptic plasticity and impaired spatial cognition in mice transgenic for exon 1 of the human Huntington’s disease mutation. J. Neurosci. 20, 5115–23 (2000).

42. Vonsattel, J.-P. et al. Neuropathological Classification of Huntinqton’s Disease. J. Neuropathol. Exp. Neurol. 44, 559–577 (1985).

43. Kelley, J. B. et al. The Defective Nuclear Lamina in Hutchinson-Gilford Progeria Syndrome Disrupts the Nucleocytoplasmic Ran Gradient and Inhibits Nuclear Localization of Ubc9. Mol. Cell. Biol. 31, 3378–3395 (2011).

44. Ferri, G., Storti, B. & Bizzarri, R. Nucleocytoplasmic transport in cells with progerin-induced defective nuclear lamina. Biophys. Chem. 229, 77–83 (2017).

45. Eftekharzadeh, B. et al. Tau Protein Disrupts Nucleocytoplasmic Transport in Alzheimer’s Disease. Neuron 99, 925–940.e7 (2018).

46. Chalovich, E. M., Zhu, J. H., Caltagarone, J., Bowser, R. & Chu, C. T. Functional repression of cAMP response element in 6-hydroxydopamine-treated neuronal cells. J. Biol. Chem. 281, 17870–17881 (2006).

47. Morton, A. J., Lagan, M. A., Skepper, J. N. & Dunnett, S. B. Progressive formation of inclusions in the striatum and hippocampus of mice transgenic for the human Huntington’s disease mutation. J. Neurocytol. 29, 679–702 (2000).

48. Hansson, O. et al. Resistance to NMDA toxicity correlates with appearance of nuclear inclusions, behavioural deficits and changes in calcium homeostasis in mice transgenic for exon 1 of the huntington gene. Eur. J. Neurosci. 14, 1492–1504 (2001).

49. Heng, M. Y. et al. Early autophagic response in a novel knock-in model of Huntington disease. Hum. Mol. Genet. 19, 3702–3720 (2010).

50. Merienne, N. et al. Cell-Type-Specific Gene Expression Profiling in Adult Mouse Brain Reveals Normal and Disease-State Signatures. Cell Rep. 26, 2477–2493.e9 (2019).

51. Park, M. et al. Age-associated chromatin relaxation is enhanced in Huntington’s disease mice. Aging (Albany. NY). 9, 803–822 (2017).

52. Valor, L. M., Guiretti, D., Lopez-Atalaya, J. P. & Barco, A. Genomic landscape of transcriptional and epigenetic dysregulation in early onset polyglutamine disease. J. Neurosci. 33, 10471–10482 (2013).

53. Bibb, J. A. et al. Severe deficiencies in dopamine signaling in presymptomatic Huntington’s disease mice. Proc. Natl. Acad. Sci. 97, 6809–6814 (2000).

54. Dash, S. K. et al. Self-assembled betulinic acid protects doxorubicin induced apoptosis followed by reduction of ROS-TNF-$α$-caspase-3 activity. Biomed. Pharmacother. 72, 144–157 (2015).

55. Saavedra, A. et al. Striatal-enriched protein tyrosine phosphatase expression and activity in huntington’s disease: A STEP in the resistance to excitotoxicity. J. Neurosci. 31, 8150–8162 (2011).

56. Arlotta, P. et al. Neuronal subtype-specific genes that control corticospinal motor neuron development in vivo. Neuron 45, 207–221 (2005).

57. Bagri, A. et al. The chemokine SDF1 regulates migration of dentate granule cells. Development 129, 4249–4260 (2002).

58. Arlotta, P., Molyneaux, B. J., Jabaudon, D., Yoshida, Y. & Macklis, J. D. Ctip2 controls the differentiation of medium spiny neurons and the establishment of the cellular architecture of the striatum. J. Neurosci. 28, 622–632 (2008).

59. Herculano-Houzel, S. & Lent, R. Isotropic fractionator: A simple, rapid method for the quantification of total cell and neuron numbers in the brain. J. Neurosci. 25, 2518–2521 (2005).

60. Alvarez-Periel, E. et al. Cdk5 Contributes to Huntington’s Disease Learning and Memory Deficits via Modulation of Brain Region-Specific Substrates. Mol. Neurobiol. 55, 6250–6268 (2018).

61. Sadaie, M. et al. Redistribution of the Lamin B1 genomic binding profile affects rearrangement of heterochromatic domains and SAHF formation during senescence. Genes Dev. 27, 1800–1808 (2013).

62. Buenrostro, J. D., Giresi, P. G., Zaba, L. C., Chang, H. Y. & Greenleaf, W. J. Transposition of native chromatin for fast and sensitive epigenomic profiling of open chromatin, DNA-binding proteins and nucleosome position. Nat. Methods 10, 1213–1218 (2013).

63. Langmead, B., Trapnell, C., Pop, M. & Salzberg, S. L. Ultrafast and memory-efficient alignment of short DNA sequences to the human genome. Genome Biol. 10, R25 (2009).

64. Kent, J. W. et al. The human genome browser at UCSC. Genome Res. 12, 996–1006 (2002).

65. Trapnell, C., Pachter, L. & Salzberg, S. L. TopHat: Discovering splice junctions with RNA-Seq. Bioinformatics 25, 1105–1111 (2009).

66. Love, M. I., Huber, W. & Anders, S. Moderated estimation of fold change and dispersion for RNA-seq data with DESeq2. Genome Biol. 15, 1–21 (2014).

67. Zhang, Y. et al. Model-based analysis of ChIP-Seq (MACS). Genome Biol. 9, R137 (2008).

68. O’Leary, N. A. et al. Reference sequence (RefSeq) database at NCBI: Current status, taxonomic expansion, and functional annotation. Nucleic Acids Res. 44, D733–D745 (2016).

69. Ye, T. et al. seqMINER: An integrated ChIP-seq data interpretation platform. Nucleic Acids Res. 39, 1–10 (2011).

70. Bailey, T. L. et al. MEME Suite: Tools for motif discovery and searching. Nucleic Acids Res. 37, 202–208 (2009).

71. Huang, D. W., Sherman, B. T. & Lempicki, R. A. Systematic and integrative analysis of large gene lists using DAVID bioinformatics resources. Nat. Protoc. 4, 44–57 (2009).

72. Barrett, T. et al. NCBI GEO: Archive for functional genomics data sets - Update. Nucleic Acids Res. 41, 991–995 (2013).

73. Ding, X. et al. Activity-induced histone modifications govern Neurexin-1 mRNA splicing and memory preservation. Nat. Neurosci. 20, 690–699 (2017).

74. Mews, P. et al. Acetyl-CoA synthetase regulates histone acetylation and hippocampal memory. Nature 546, 381–386 (2017).

75. Sams, D. S. et al. Neuronal CTCF Is Necessary for Basal and Experience-Dependent Gene Regulation, Memory Formation, and Genomic Structure of BDNF and Arc. Cell Rep. 17, 2418–2430 (2016).

