## Supplementary Data for "Neuron-specific increase in lamin B1 disrupts nuclear function in Huntington’s disease"

Supplementary Figure 1

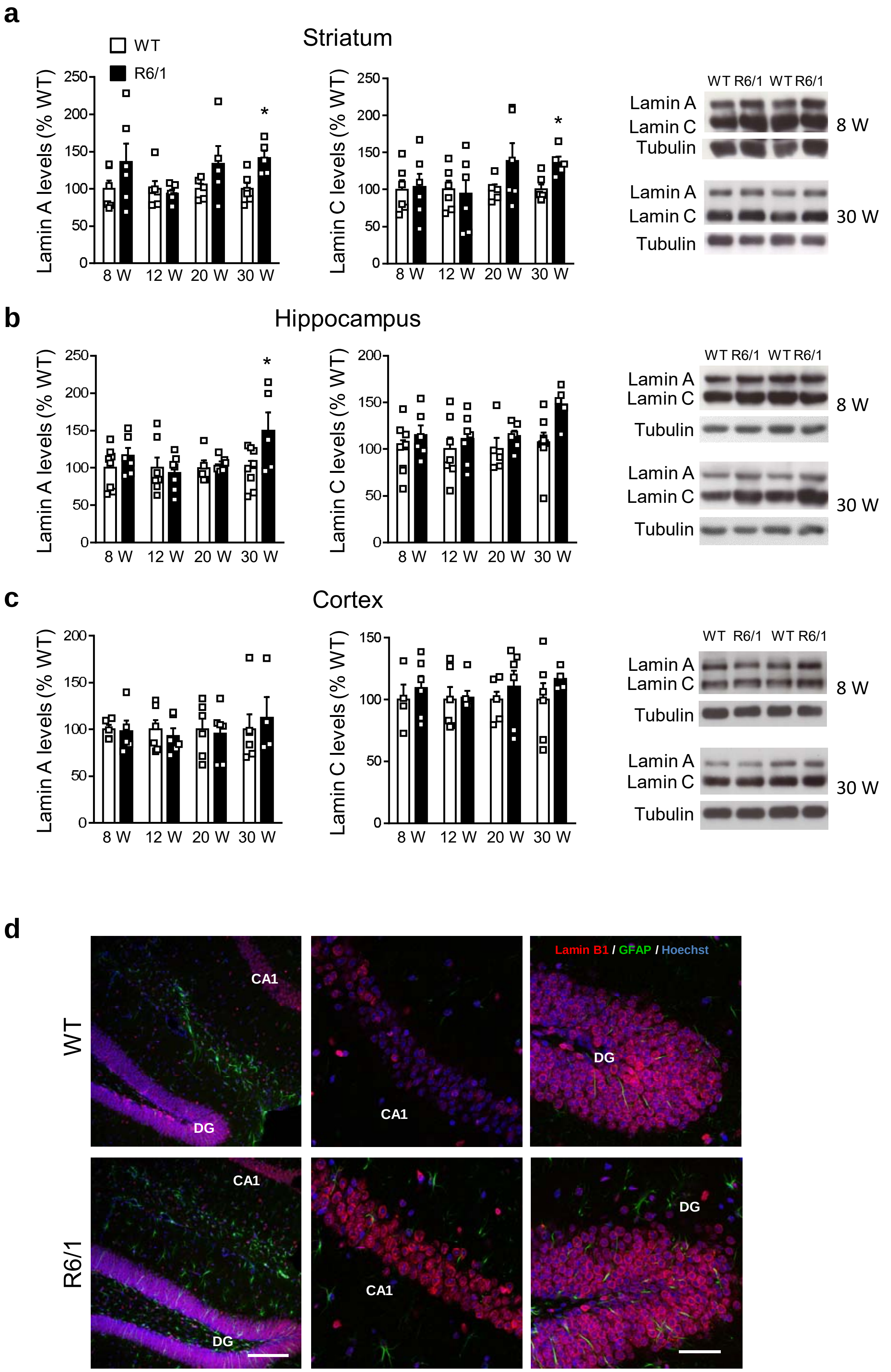

**Supplementary Fig. 1 Lamin A and C protein levels in different mouse brain regions at different ages, and lamin B1 localization in mouse hippocampus at 12 weeks of age.** Lamin A and lamin C protein levels in the striatum (**a**), hippocampus (**b)** and cortex (**c)** of wild-type (WT) and R6/1 mice at different stages of the disease progression (W: weeks). Each point corresponds to the value from an individual sample. Bars represent the mean ± S.E.M. Two-tailed unpaired Student's t test. *$p < 0.05$ as compared with WT mice. Representative immunoblots are shown. **d,** Representative images showing the distribution of lamin B1 in the hippocampus of wild-type (WT) and R6/1 mice at 12 weeks of age. Antibody against lamin B1 (red) was combined with anti-GFAP antibody (green) to label astrocytes and Hoechst 33258 (blue) to label nuclei. DG: Dentate gyrus.

### Supplementary Figure 2

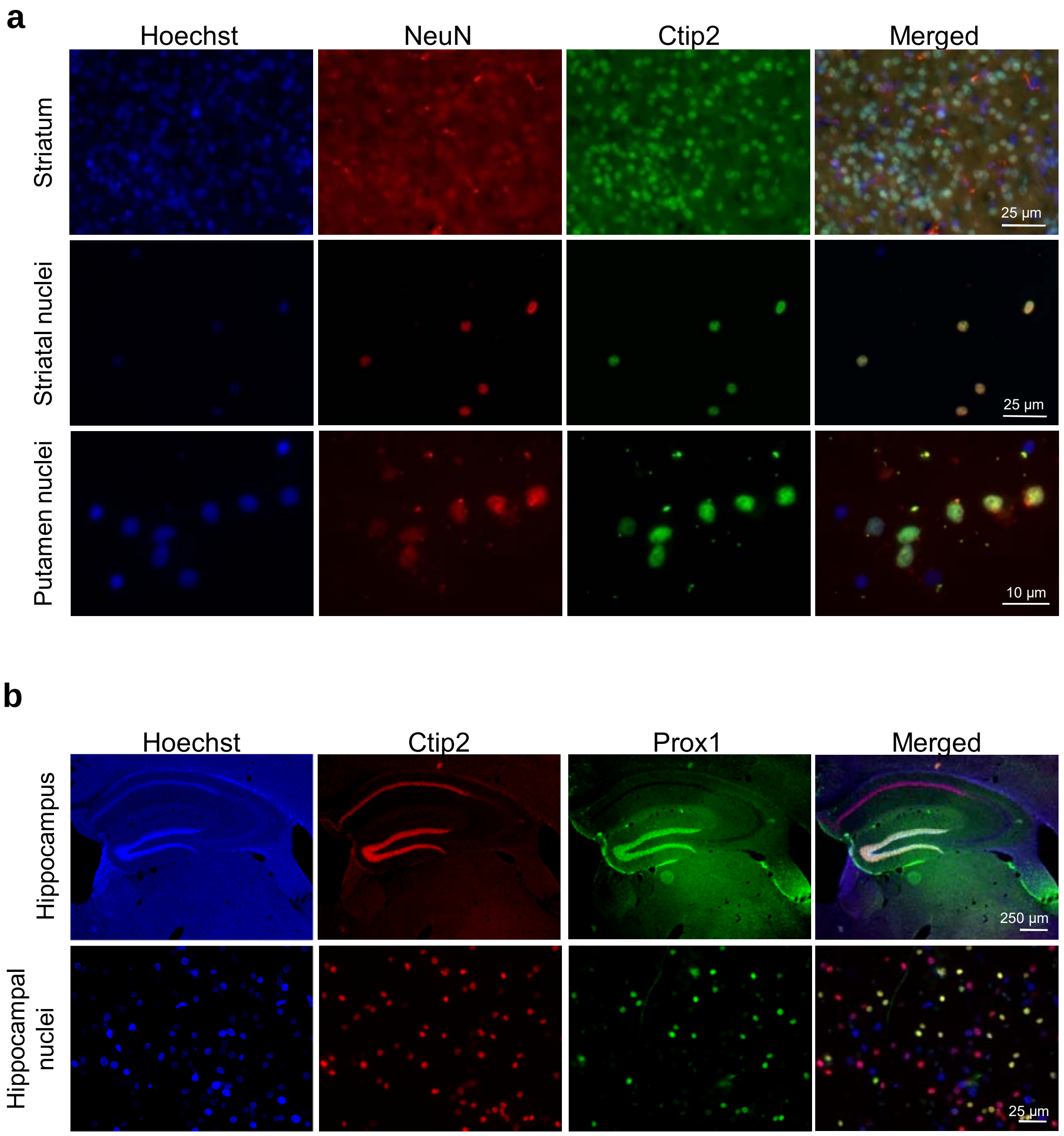


**Supplementary Fig. 2 Nuclear markers for striatal MSNs and hippocampal CA1 and DG neurons. a,** Superior panels show a striatal slice stained with Hoechst 33258 and antibodies used for nuclei selection. Images below show the correlative staining of the upper marker after the nuclear purification in R6/1 mice striatum (striatal nuclei) and in the putamen of HD patients (Putamen nuclei). The right panels represent the merge of the three other images. Classification of the nuclei can be clearly performed according to their colour being MSNs nuclei the yellow ones. **b,** Superior panels show a hippocampal slice stained with Hoechst 33258 and antibodies used for nuclei selection according to their localization in the original tissue. The panels below show the correlative staining of the upper marker after the nuclear purification. The right panels represent the merge of the three other images. Classification of the nuclei can be clearly performed according to their colour (yellow for DG neurons nuclei and magenta for CA1 neurons nuclei).

Supplementary Figure 3

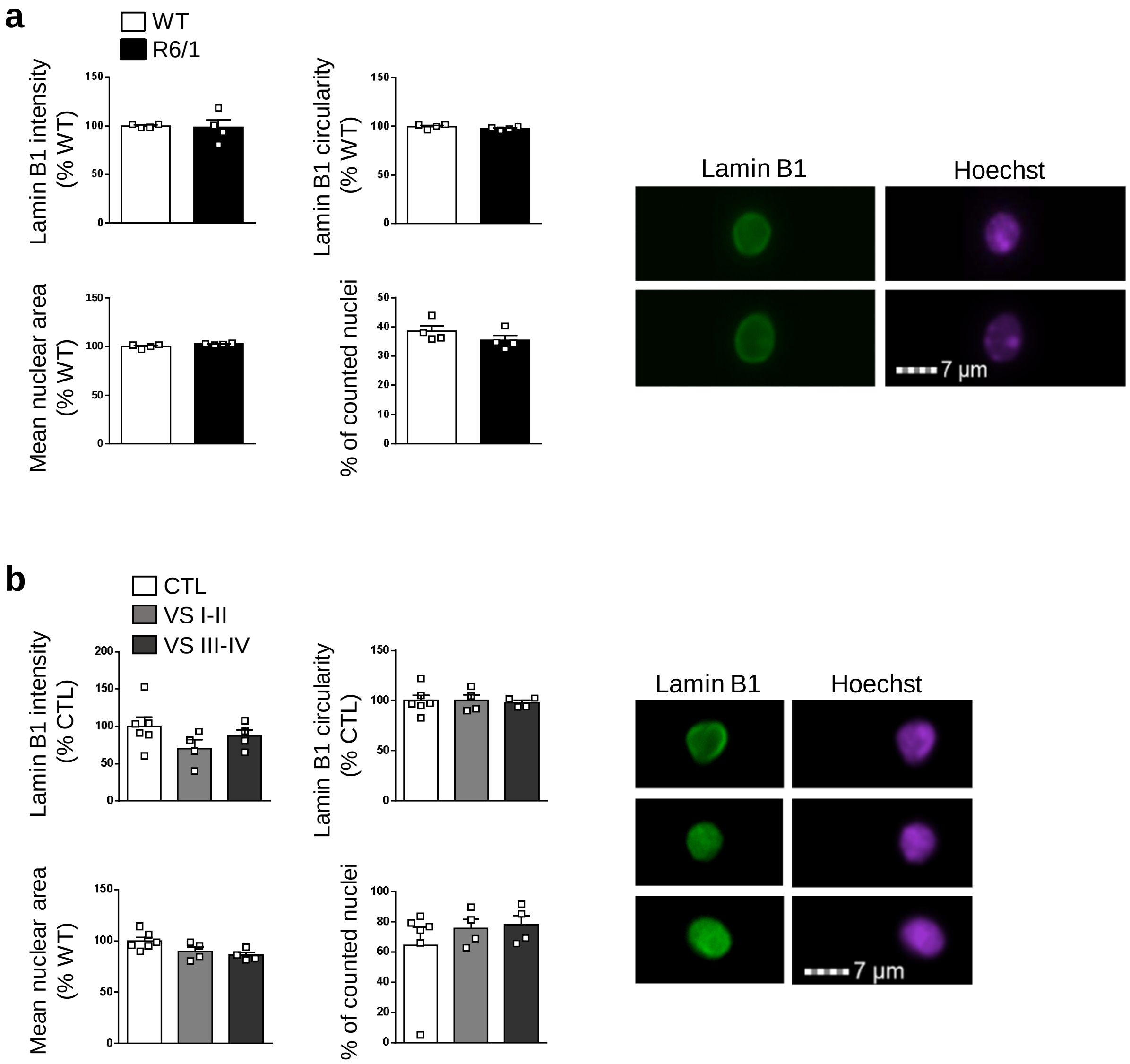


**Supplementary Fig. 3 Lamin B1 levels and nuclear morphology are not altered in striatal glial cells nuclei. a,** Graphs show the quantification of different parameters in striatal glial nuclei (lamin B1+/Hoescht+) from 30-week-old wild-type (WT) and R6/1 mice. Representative images are shown. **b,** Graphs show the quantification of different parameters in striatal glial nuclei from HD patients at different stages of the disease (Vonsattel grades I-II and III-IV). Representative images are shown. In graphs, each point corresponds to the value from an individual sample. Bars represent the mean ± S.E.M.

### Supplementary Figure 4

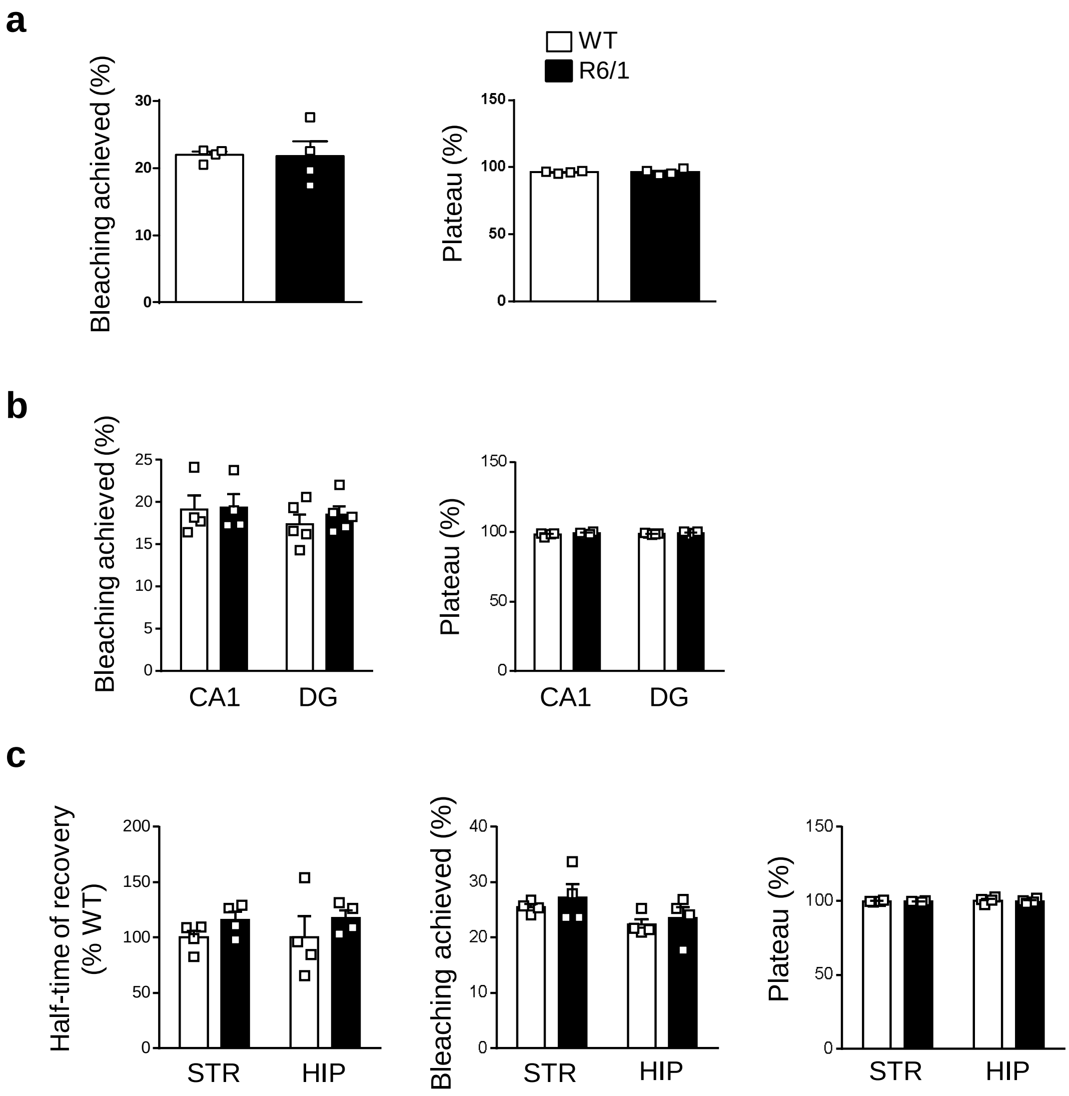


**Supplementary Fig. 4 Dextran freely penetrates in striatal and hippocampal neurons nuclei. a,** Graphs show the quantification of different parameters in MSNs nuclei from 30-week old wild-type (WT) and R6/1 mice in the FRAP experiment. **b,** Graphs show the quantification of different parameters in hippocampal CA1 and DG nuclei from 30-week old wild-type (WT) and R6/1 mice in the FRAP experiment. **c** FRAP experiment in the background showed no differences in any of the parameters analyzed

### Supplementary Figure 5

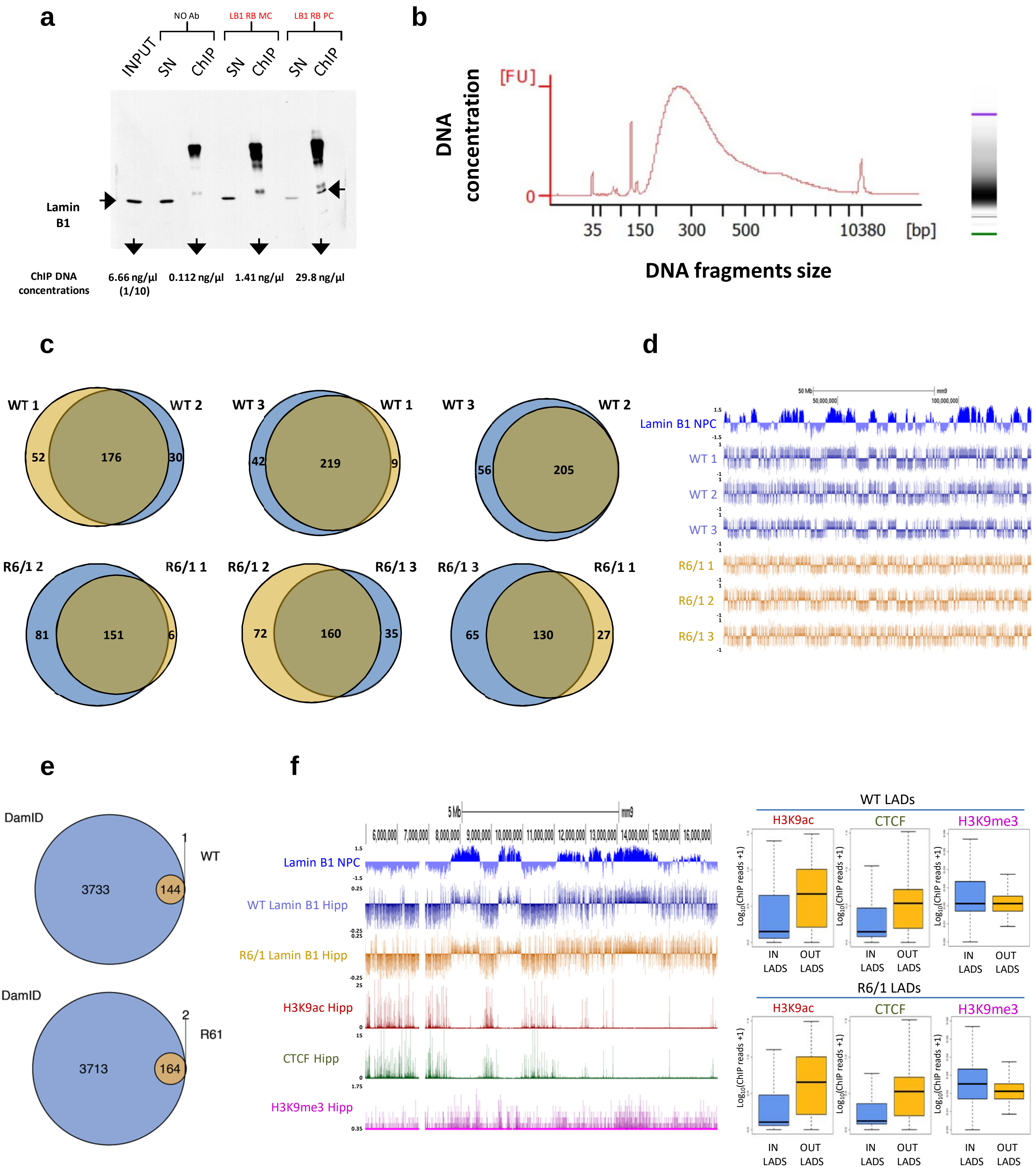

**Supplementary Fig. 5 Lamin B1 ChIP-seq and nuclear sub-fractionation additional data**. **a,** Western-blot of immunoprecipitated fractions obtained from lamin B1 ChIp-seq. Lamin B1 protein levels were analysed in Input (total DNA fraction), supernatant (SN) and chromatin immunoprecipitated (ChIP) fractions in no antibody conditions (No ab), using a mouse monoclonal antibody against lamin B1 (LB1 MS MC, ab8982) and using a rabbit polyclonal antibody against lamin B1 (LB1 RB PC, ab16048). DNA concentrations of eluted DNA from each fraction is shown. **b,** Bioanalyzer profile of sonicated chromatin from ChIP-seq experiment. **c,** Venn diagrams showing the number of overlapping LADs identified by EDD among the different generated replicates for WT and R6/1 mice datasets. **d,** UCSC genome browser capture of lamin B1 NPC DamID data and ChIP-seq signal (log(LB1/Input)) for all replicates generated using WT and R6/1 mice hippocampal tissue. **e,** Venn diagrams showing the number of overlapping LADs between previous DamID characterized NPCs LADs and WT (top) or R6/1 (bottom) mice hippocampus identified LADs. **f,** UCSC genome browser capture of lamin B1 NPC DamID data, WT and R6/1 mice hippocampus ChIP-seq signal (log(LB1/Input)) and publically available data from WT mice hippocampus of H3K9ac, CTCF and H3K9me3 ChIP-seq data (left). H3K9ac, CTCF and H3K9me3 ChIP-seq data enrichment in WT (right top) and R6/1 (right bottom) mice hippocampus LADs.

### Supplementary Figure 6

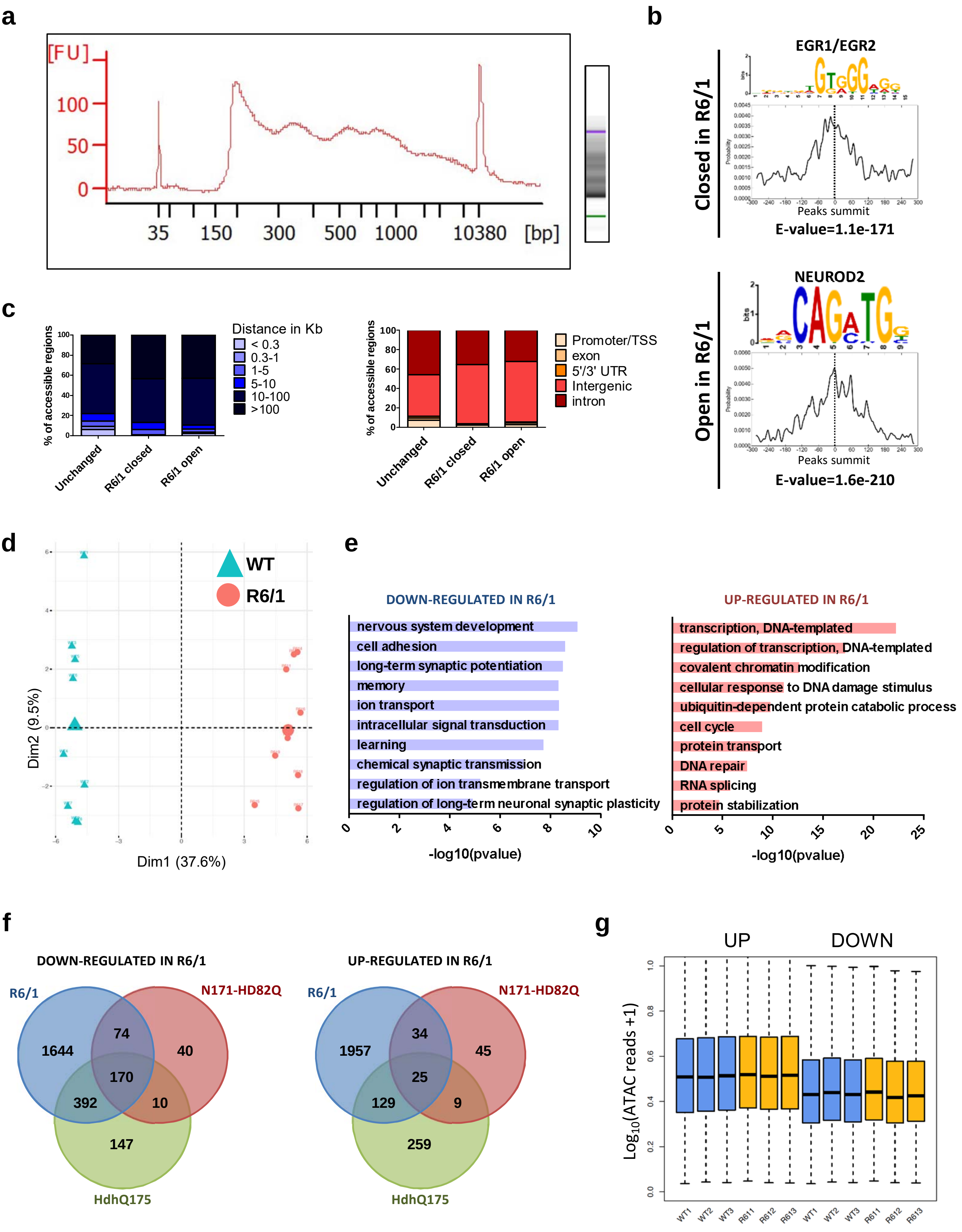

**Supplementary Fig. 6 ATAC-seq and RNA-seq additional data. a,** Bioanalyzer profile of transposed DNA from ATAC-seq experiments. **b,** Motif analysis of closed (top) and opened (bottom) chromatin regions in R6/1 mice. Associated transcription factor, motif distribution and E-value are shown for each discovered motif. **c,** Stacked bars graphs showing unchanged, R6/1 closed and R6/1 open chromatin regions distance to closest TSS (left) and distribution among annotated genomic categories (right). **d,** Principal component analysis (PCA) of RNA-seq data generated in WT and R6/1 mice hippocampus, N=9. **e,** Bargraphs of significiant (Benjamini's adjuted pvalue < 0.05) Biological Processes (BP) terms from DAVID for genes down- (left) and up-regulated (right) in R6/1 mice hippocampus (adjpvalue <0.001). Bars represents the $-\log_{10}$(Benjamini's adj pvalue). **f,** Venn diagrams showing the overlap of down- (left) and up-regulated (right) genes in R6/1 mice hippocampus (adjpvalue <0.001) with N171-HD82Q and HdhQ175 differentially expressed genes previously identified. **g,** Boxplots showing average TSS chromatin accessibility ($\log_{10}$(ATAC reads +1), N=3) for genes up or down-regulated (adjpvalue<0.001, N=9) in R6/1 mice hippocampus (right).

### Supplementary Figure 7

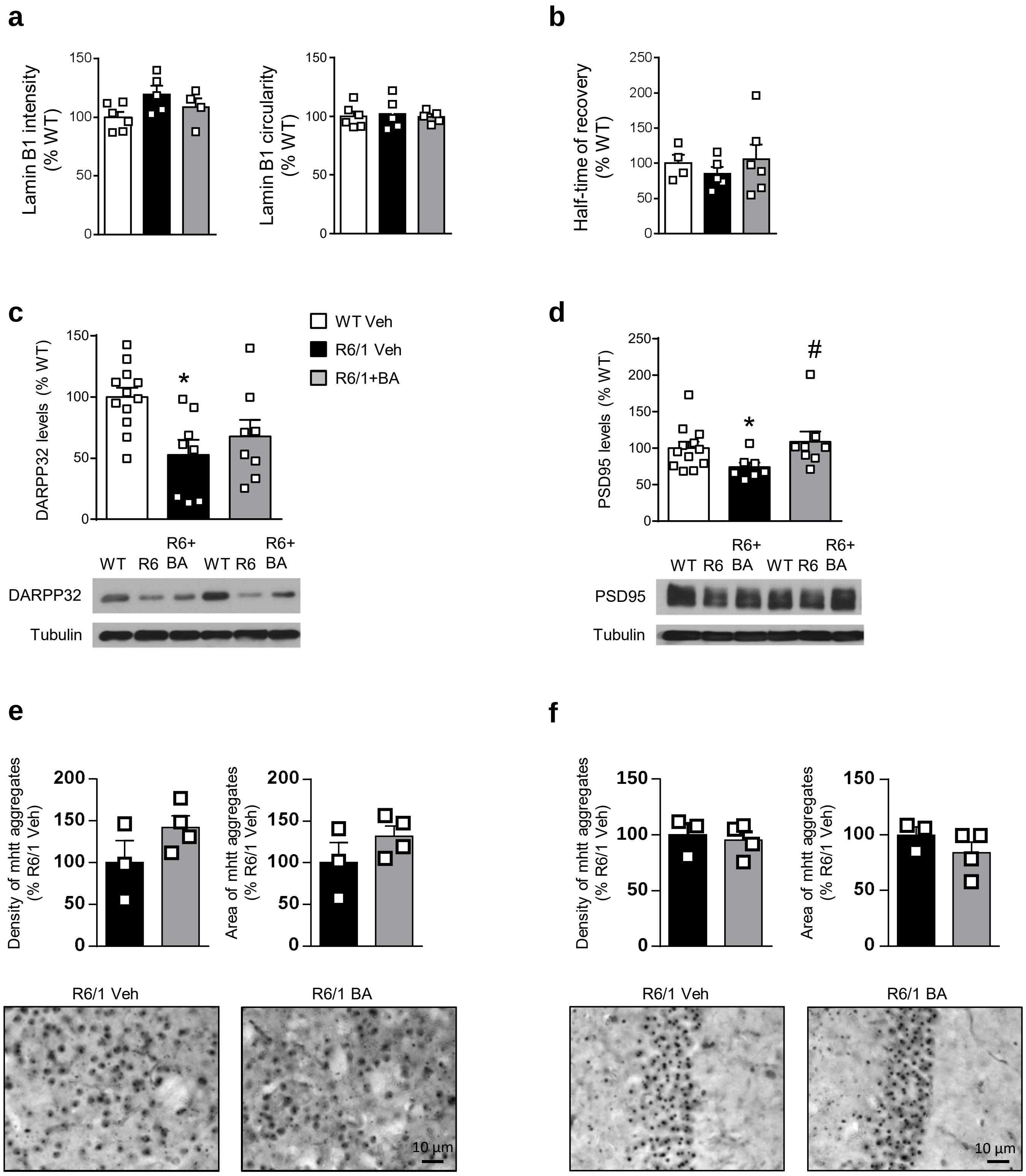

**Supplementary Fig. 7 Effects of betulinic acid in the striatum and hippocampus at the molecular level. a,** Lamin B1 intensity and circularity in the nucleus of hippocampal DG neurons (Ctip2+/Prox1+). Graphs show the quantification in wild-type (WT) and R6/1 mice 12 weeks after treatment (Veh, vehicle; BA, betulinic acid). **b,** Nuclear permeability in hippocampal DG neurons nuclei from wild-type (WT) and R6/1 mice 12 weeks after treatment. **c,** Striatal DARPP32 and **d**, hippocampal PSD95 levels in wild-type (WT) and R6/1 mice 12 weeks after treatment (veh, vehicle; BA, betulinic acid). $P < 0.05$ compared to vehicle-treated WT mice (one-way ANOVA followed by Bonferroni's *post hoc* test). **e-f,** Brain slices from R6/1 mice were processed for immunohistochemistry using the EM48 antibody to detect mhtt aggregates in **e,** the striatum and **f,** hippocampus. The number and area of mhtt aggregates in each group of mice were evaluated using stereological tools. Representative images are shown. **a-f,** Each point corresponds to the value from an individual sample. Bars represent the mean ± S.E.M.

Supplementary Figure 8

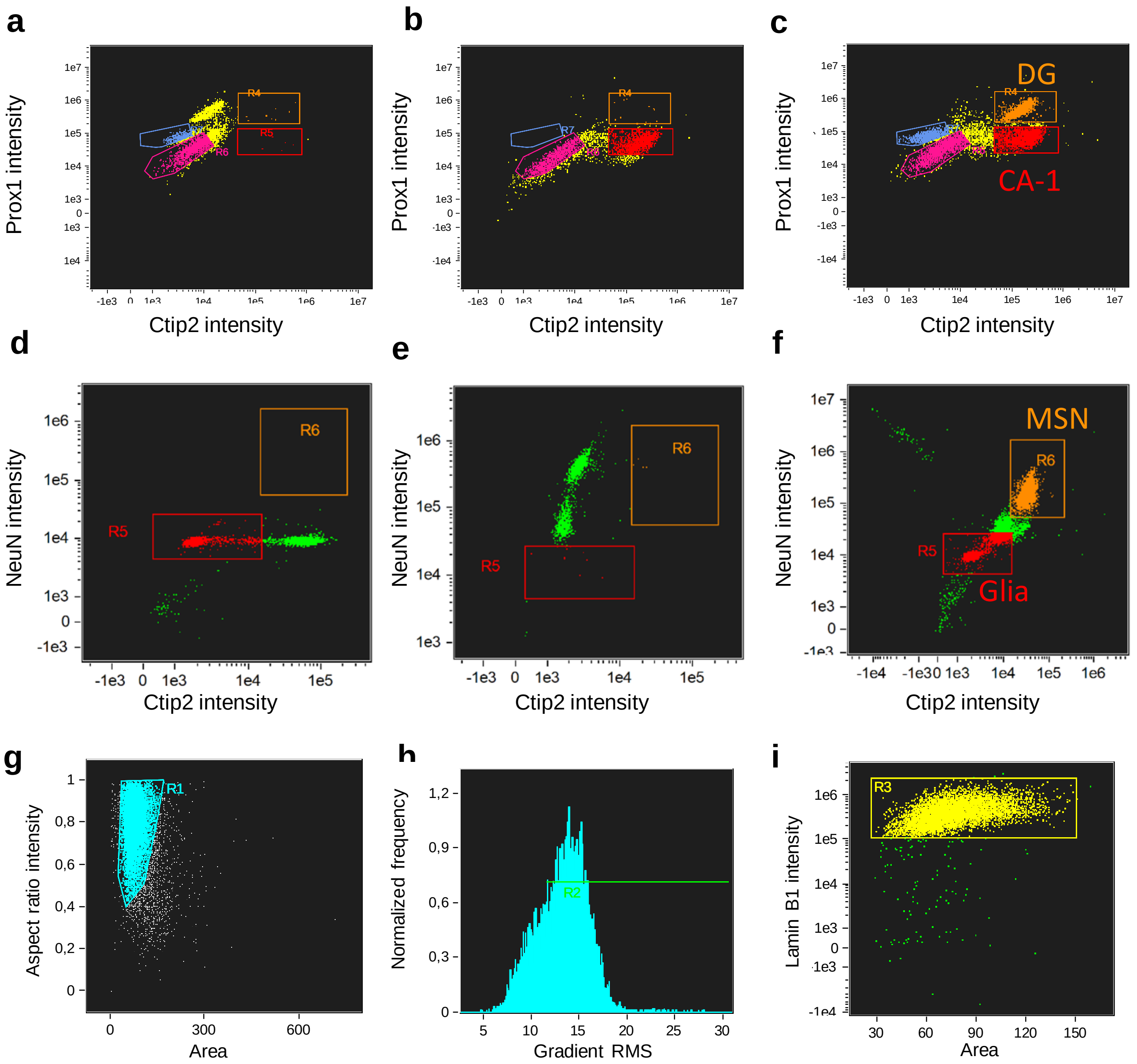


**Supplementary Fig. 8 Selection of nuclei from neuronal populations**. Identification of different populations in the (**a-c**) hipocampus and (**d-f**) striatum using FMO for the different antibodies (**a**: Ctip2; **b**: Prox1; **d**: double FMO for Ctip2 and NeuN; **e**: combination of single FMO for Ctip2 and NeuN). The final populations analyzed are represented (**c, f**). **g,** Representation of all the recorded events according to their area (x axis) and aspect ratio (y axis). The singlets population (individual nuclei) was selected according to the most condensed area of nuclei with an aspect ratio over 0.4 and an area of around 50-100 µm. **h,** Internal parameter used as a measure of focused images (gradient RMS), and only events with a gradient RMS value over 14 were accepted as correctly focused images. **i,** Focused events graphed according to their area and lamin B1 intensity. Only events clustered in the dense cloud were considered as real nuclei, discarding possible debris present in the sample.

### Supplementary Figure 9

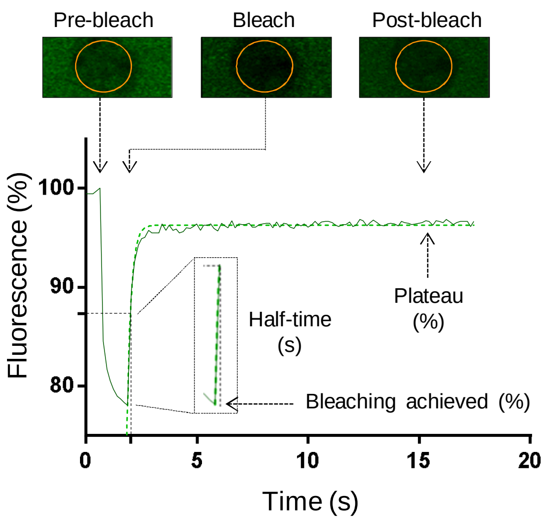


**Supplementary Fig. 9 Representation of FRAP recovery curve with the parameters analyzed.** Representative images of pre-bleach, bleach and post-bleach phases are shown. Orange circle: ROI

**Supplementary Table 1 Details on human *post-mortem* brain samples**

| ID | Pathological diagnosis | Gender | Age (years) | CAG repeats | PMD (hh:mm) |
|---|---|---|---|---|---|
| BK-0810 | Normal | Female | 81 | - | 23:30 |
| BK-1570 | Normal | Female | 86 | - | 4:00 |
| BK-1679 | Normal | Female | 90 | - | - |
| BK-1774 | Normal | Female | 74 | - | - |
| BK-1491 | Normal | Male | 83 | - | 13:00 |
| BK-1563 | Normal | Male | 79 | - | - |
| BK-1697 | Normal | Male | 78 | - | 6:00 |
| BK-1733 | Normal | Male | 76 | - | 11:30 |
| CS-1334 | HD, Vonsattel grade 1 | Male | 73 | 40 +/-2 | 7:00 |
| CS-1630 | HD, Vonsattel grade 2 | Male | 76 | 41 | 6:00 |
| CS-1638 | HD, Vonsattel grade 2 | Male | 72 | - | 13:10 |
| CS-1758 | HD, Vonsattel grade 2-3 | Male | 68 | 42 +/-2 | 6:10 |
| CS-1438 | HD, Vonsattel grade 3 | Male | 85 | 40 | 5:30 |
| CS-1120 | HD, Vonsattel grade 3 | Male | 55 | 48 | 15:00 |
| CS-1193 | HD, Vonsattel grade 3-4 | Male | 55 | - | 7:00 |
| CS-1294 | HD, Vonsattel grade 3 | Male | 53 | 45 +/-2 | 7:00 |

Information about the gender, age, CAG repeat length, Vonsattel grade and *post-mortem* delay (PMD) from the 15 control individuals and 12 HD patients analyzed.

#### Supplementary Table 2 Details on antibodies used in the study

| ANTIBODY | SOURCE | IDENTIFIER | DILUTION |
|---|---|---|---|
| **WESTERN BLOT** | | | |
| Rabbit anti-Lamin B1 polyclonal antibody (human samples) | Abcam | Cat# ab16048, RRID:AB_443298 | 1:1000 |
| Mouse anti-Lamin B1 monoclonal antibody (mouse samples) | Abcam | Cat# ab8982, RRID:AB_1640627 | 1:1000 |
| Mouse anti-Lamin B2 | Santa Cruz Biotechnology | Cat# ab18465, RRID:AB_2064130 | 1:1000 |
| Rabbit anti-Lamin A/C (H-110) | Santa Cruz Biotechnology | Cat# sc-20681, RRID:AB_648154 | 1:1000 |
| Mouse anti-human TATA binding protein (TBP) | Abcam | Cat# ab51841, RRID:AB_945758 | 1:1000 |
| Mouse anti-PSD95 (7E3-1B8) | Thermo Fisher Scientific | Cat# MA1-046, RRID:AB_2092361 | 1:1000 |
| Mouse anti-DARPP32, clone 15 | BD Bioscience | Cat# 611520; RRID: AB_398980 | 1:1000 |
| Mouse anti-tubulin | Sigma-Aldrich | Cat# T5168; RRID: AB_477579 | 1:50,000 |
| Anti-Rabbit IgG (H+L), HRP Conjugate | Promega | Cat# W4011, RRID:AB_430833 | 1:2000 |
| Anti-Mouse IgG (H+L), HRP Conjugate | Promega | Cat# W4021, RRID:AB_430834 | 1:2000 |
| **IMMUNOHISTOCHEMISTRY** | | | |
| Mouse anti-Glial fibrillary acidic protein | Sigma-Aldrich | Cat# G3893, RRID:AB_477010 | 1.200 |
| Rabbit anti-Lamin B1 polyclonal antibody (human samples) | Abcam | Cat# ab16048, RRID:AB_443298 | 1:150 (mouse sections); 1:50 (human sections) |

| | | | |
|---|---|---|---|
| Mouse anti-Huntingtin, clone EM48 | Millipore | Cat# mab5374; RRID: AB_10055116 | 1:150 |
| Cy3-AffiniPure F(ab')2 Fragment Goat Anti-Rabbit IgG, F(ab')2 Fragment Specific | Jackson ImmunoResearch Labs | Cat# 111-166-047, RRID:AB_2338010 | 1:200 |
| Cy2-AffiniPure Goat Anti-Mouse IgG (H+L) | Jackson ImmunoResearch Labs | Cat# 115-225-146, RRID:AB_2307343 | 1:200 |
| **ISOLATED NUCLEI** | | | |
| Rabbit anti-Lamin B1 polyclonal antibody (human samples) | Abcam | Cat# ab16048, RRID:AB_443298 | 1:400 (hippocampus); 1:800 (striatum) |
| Rat anti-Ctip-2 (25B6) | Abcam | Cat# ab18465, RRID:AB_2064130 | 1:400 |
| Mouse anti-Prox1, clone 5G10 | Millipore | Cat# MAB5652, RRID:AB_827462 | 1:200 |
| Mouse anti-NeuN | Millipore | Cat# MAB377, RRID:AB_2298772 | 1:800 |
| Alexa Fluor® 488-AffiniPure Donkey Anti-Rabbit IgG (H+L) | Jackson ImmunoResearch Labs | Cat# 711-545-152, RRID:AB_2313584 | 1:250 |
| Alexa Fluor 647-AffiniPure Goat Anti-Mouse IgG (H+L) | Jackson ImmunoResearch Labs | Cat# 115-605-166, RRID:AB_2338914 | 1:250 |
| Donkey F(ab')2 Anti-Rat IgG H&L (Alexa Fluor® 555) preadsorbed | Abcam | Cat# ab150150 | 1:250 |
